# Gene networks reveal stem-cell state convergence during preneoplasia and progression to malignancy in multistage skin carcinogenesis

**DOI:** 10.1101/2023.05.08.539863

**Authors:** Mark A. Taylor, Eve Kandyba, Kyle Halliwill, Reyno Delrosario, Matvei Koroshkin, Hani Goodarzi, David Quigley, Yun Rose Li, Di Wu, Saumya Bollam, Olga Mirzoeva, Rosemary J. Akhurst, Allan Balmain

## Abstract

Adult mammalian stem cells play critical roles in normal tissue homeostasis, as well as in tumor development, by contributing to cell heterogeneity, plasticity, and development of drug resistance. The relationship between different types of normal and cancer stem cells is highly controversial and poorly understood. Here, we carried out gene expression network analysis of normal and tumor samples from genetically heterogeneous mice to create network metagenes for visualization of stem-cell networks, rather than individual stem-cell markers, at the single-cell level during multistage carcinogenesis. We combined this approach with lineage tracing and single-cell RNASeq of stem cells and their progeny, identifying a previously unrecognized hierarchy in which *Lgr6*+ stem cells from tumors generate progeny that express a range of other stem-cell markers including *Sox2, Pitx1, Foxa1, Klf5*, and *Cd44*. Our data identify a convergence of multiple stem-cell and tumor-suppressor pathways in benign tumor cells expressing markers of lineage plasticity and oxidative stress. This same single-cell population expresses network metagenes corresponding to markers of cancer drug resistance in human tumors of the skin, lung and prostate. Treatment of mouse squamous carcinomas *in vivo* with the chemotherapeutic *cis*-platin resulted in elevated expression of the genes that mark this cell population. Our data have allowed us to create a simplified model of multistage carcinogenesis that identifies distinct stem-cell states at different stages of tumor progression, thereby identifying networks involved in lineage plasticity, drug resistance, and immune surveillance, providing a rich source of potential targets for cancer therapy.

**One-Sentence Summary:** Genes act in networks to drive cancer, and we identify these groups of genes from bulk-tissue and trace them at single-cell resolution.

## Main Text

### The skin as a model for normal and cancer stem-cell analysis

The development of phenotypic plasticity is a common, and probably universal, feature of cancer development that was recently recognized as an emerging cancer hallmark(*1*). The mechanistic basis for this plasticity, defined as the redeployment of lineage-specific gene expression programs along alternative cell fate trajectories, is presently unclear (*2*). Resolution of these questions has major practical implications for understanding and treatment of cancer, since cell plasticity has been associated with invasion, metastasis, and resistance to chemotherapy or targeted drugs(*3*–*5*). Tumor cell plasticity has been attributed to the existence of cancer stem cells (CSCs), but there is presently no consensus as to how CSCs relate to their normal tissue counterparts, to the cells of origin of tumors, or even whether they exist(*6, 7*). Different models suggest that CSCs may lie within a hierarchy of differentiation within tumors (*8, 9*) or they may show complete plasticity, being essentially interconvertible(*10*).

Here, we address these questions using mouse skin, which has over several decades been the most widely studied solid tissue for analysis of stem-cell function in normal homeostasis and during oncogenic transformation(*11*–*13*). In the mouse, a number of distinct stem-cell markers have been identified which contribute to homeostasis by repopulating specific compartments within the epithelium(*14, 15*). Certain stem cells in normal skin have also been proposed to act as cancer stem cells (CSCs) in several tumor types(*9, 16*), marked by *Lgr5*(*17*), *Lgr6*(*18*), *Twist1* (*19*), *Sox2*(*20*), and *Pitx1*(*21*). To identify the cell plasticity states that arise during transitions from normal tissue through pre-neoplasia to malignant carcinomas, we developed a model system that encompasses both germline genetic and somatic genetic diversity to construct gene expression network modules representing each stage, and visualized these networks in single cells representing the continuum of steps in carcinogenesis. Our data identify previously unappreciated connections between different stem cell populations in the skin, leading to a more simplified model of cell state transitions during tumorigenesis.

### Rewiring of stem-cell gene expression networks in tumors

Although progress has been made in unravelling gene networks in single cells(*22*), understanding global gene network dysregulation in cancer continues to rely upon bulk-tissue data(*23*). Bulk-tissue data are useful because gene networks are generated from hundreds of independent samples and thus are likely general features of this system, occurring repeatedly across separate instantiations of carcinogenesis. On the other hand, features in single cells are usually inferred from a small number of samples because of the logistical trade-off between the number of samples and the number of cells that ‘-omics’ assays can measure. Thus single-cell assays represent specific contingent outcomes of a particular sample rather than general system features(*24*). Together, the sampling breadth and cellular resolution of bulk-tissue and single-cell modalities are complementary. By identifying coordinated gene expression changes that are common across hundreds of bulk samples, we can identify conserved features of carcinogenesis. By then combining this with stem-cell biology at the single-cell level, we observe how these conserved features operate in single cells to drive initiated cells from normal homeostasis to cancer.

We previously carried out genetic and gene expression analysis of multistage carcinogenesis, resulting in transcriptomic profiles of normal skin, and chemically-induced benign, malignant, and metastatic carcinomas from a genetically heterogeneous mouse population(*25*–*27*). Chemical carcinogenesis models capture a realistic view of cancer development in human populations since 1) tumors are induced by environmental insult, rather than genetic modification, and carry thousands of somatic mutations including many cancer driver mutations that are also seen in human tumors(*26, 28, 29*); 2) they are autochthonous to the animals in which they are generated, rather than transplanted into host animals; and 3) the treated mice are from genetically heterogeneous mouse populations that mimic human germline diversity. From this, we accumulated a transcriptomic database comprising several hundred samples that span progression from normal skin to metastasis(*25, 26*). A similar database has been used to infer functions for specific genes based on their network structures in bulk-tissue samples(*25, 27, 30, 31*). These networks are based on gene-gene correlations across samples, and thus capture indirect interactions among genes. For the present analysis, we considered indirect interactions to be biologically relevant effects of a gene’s function, so we did not seek to limit network inference to specific physical interactions such as transcription factors and their targets(*32*) or among their protein products(*33*). Here, we profiled gene expression of 106 normal skin and 157 carcinoma samples from interspecific *Mus spretus* x *FVB/N* backcross mice, generated as previously described(*34*) (Fig.1a).

**Fig. 1.**
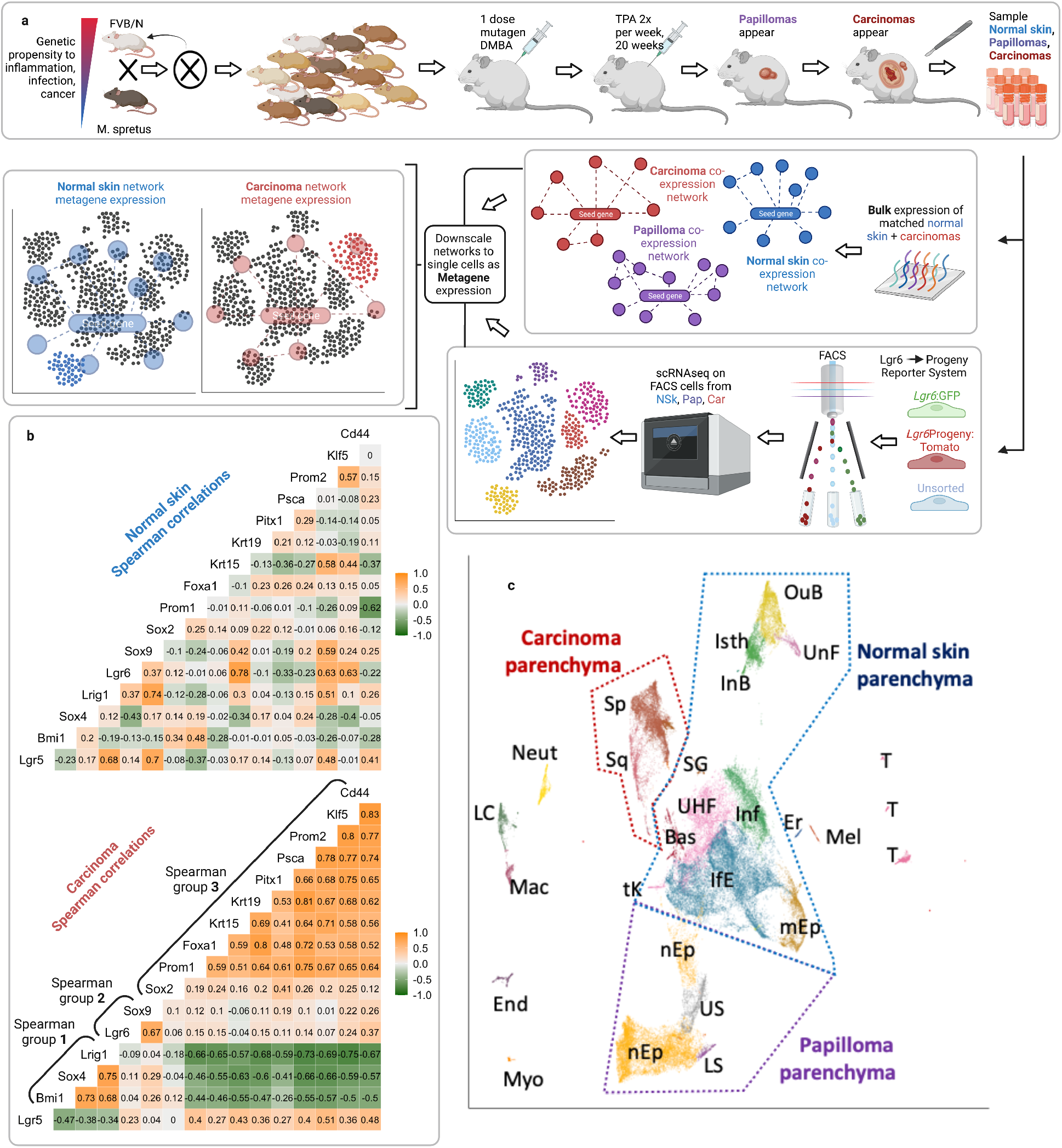
Bulk-tissue and single-cell expression data. (**a**), Experimental pipeline showing creation of a backcross mapping population segregating genetic variation in resistance to inflammation, infection, and cancer. Mice were randomly selected for subsequent matched normal skin, papilloma, and tumor expression analysis. Tumor initiation occurred by DMBA application to dorsal back skin and promotion by twice weekly TPA treatment for 20 weeks. Matched normal skin and tumors were co-sampled for bulk gene expression assays by microarray. Co-expression gene networks for genes-of-interest (GOIs) were inferred from bulk expression data. Several normal skin, papilloma and carcinoma lesions from inbred FVB mice were profiled using scRNAseq. Network expression inferred from bulk samples was overlaid onto the single-cell data as tissue-specific metagenes: a normal-skin metagene generated from normal skin bulk-tissue samples and a carcinoma metagene generated from carcinoma bulk-tissue samples. (**b)** Spearman rank correlations of selected stem cell-related genes in 106 normal skin and 157 carcinoma (Car) samples. (**c**) Single-cell data from normal skin, papilloma and carcinoma from FVB mice displayed on a UMAP with cells colored according to type; OuB is for outer bulge; Isth isthmus; UnF undifferentiated follicular; InB inner bulge; SG sebaceous gland; UHF upper hair follicle; Bas basal cells; tK terminal keratinocytes; IfE interfollicular epidermal cells; Inf infundibulum; mEp mitotic epithelial cells; Er erthryocytes; Mel melanocytes; T T-cells; nEp neoplastic epithelia of the papilloma; LS lower spike cells (defined hereafter); US upper spike cells (defined hereafter); Sp spindle cells of the carcinoma; Sq squamous cells of the carcinoma; Neut neutrophils; LC Langerhans cells; Mac macrophages; End endothelial cells; Myo myofibroblasts.

To gain an unbiased view of the global gene network, we implemented the unsupervised inference algorithm WGCNA, which clusters genes sharing similar expression from bulk-tissue samples into gene modules(*35*) (Data S1). We then tested these modules for functional enrichment using biological process gene ontologies (GO) (Data S2). One large gene module in the carcinomas, Module 3 (N_gene_=1720), was highly enriched for genes associated with wound healing. This wound healing module included a number of known stem-cell marker genes that have been implicated in carcinogenesis, including *Sox9, Psca, Pitx1, Krt15, Krt19*, and *Lgr6. Lgr6* is of particular interest in this system since *Lgr6*+ cells are highly clonogenic in squamous carcinomas(*18*), play an important role in wound healing(*36*), and are major targets for initiation by chemical carcinogens in this model system (Kandyba et al, submitted).

We next examined the correlation structure of these stem-cell marker genes in normal tissues or tumors. This revealed three clusters of genes that were correlated with each other in tumors but not in matched normal tissues (Fig. 1b). We refer to these groups of groups of correlated stem-cell genes that are co-expressed at the bulk-tissue level as ‘Spearman groups.’ These Spearman groups reflected the gene modules identified by WGCNA (Data S3). Importantly, WGCNA identified Spearman group 1, consisting of *Bmi1, Sox4*, and *Lrig1*, as strongly anti-correlated to Spearman group 3, the latter comprising multiple known adult stem-cell markers.

Each stem-cell marker was then used as a “seed gene” to generate normal skin- and carcinoma-specific gene correlation networks (called “metagenes,” Data S4) which can capture the downstream function of specific genes in bulk-tissue samples(*25, 27, 30, 31*). The *Lgr6* metagene showed significant network rewiring between normal skin and carcinoma. In normal skin, the *Lgr6* network included of a number of epidermal lineage markers including *Krt15* (ρ=0.78) and *Klf5* (ρ=0.63), as well as *Znrf3* (ρ=0.51) and *Rnf43* (ρ=0.55), the two E3-ligases that act as R-spondin coreceptors to mediate *Wnt* signaling(*37*). However during carcinogenic rewiring, these correlations were lost, replaced by connections to *Sox9* (ρ=0.67), as well as to other important drivers of cancer phenotypes including *Tgfb1* (ρ=0.60) (table S1). *Tgfb1* is a growth factor known to play multiple roles in tumor development, acting as a growth inhibitor at early stages and switching roles to become an inducer of the epithelial-mesenchymal transition (EMT) during tumor progression(*38*). It also is a major mediator of immunosuppression that, when inhibited with blocking antibodies, improves immunotherapy outcomes in mouse models(*39, 40*). Thus, changes in the correlation network architecture of the *Lgr6* metagene during tumor development revealed biologically functional rewiring associated with progression and immune escape.

### Single-cell transcriptomes in multi-stage carcinogenesis

Since our goal was to understand how gene networks inferred from hundreds of bulk samples can be visualized in single cells, we then obtained single-cell RNA sequencing data for normal skin, benign papillomas, and malignant carcinomas, and plotted metagene expression within the resulting UMAPs. Additionally, since *Lgr6*+ cells are highly clonogenic in SCCs, we carried out lineage tracing of *Lgr6+* cells in normal skin, benign papillomas and carcinomas induced in the *Lgr6-eGFP-tdTomato* mouse strain(*18*), to separate *Lgr6*+ cells from their direct progeny. We analyzed a total of 57,807 cells of which 33,234 (57.5%) cells were normal skin; 15,280 (26.4%) were papilloma; and 9,293 (16.1%) were carcinoma (Fig.1c). We identified 21 low-resolution cell clusters by unsupervised Leiden clustering, of which cell clusters 1-4 dominated the assembly, predominantly populated by NSk (1,2); Pap (3); and Car (4), which we consider to be tissue-specific parenchymal cells (fig. S1a,b). We re-analyzed the carcinoma parenchyma alone, which showed two distinct keratinocyte subpopulations (fig. S1c) corresponding to squamous (20.8% of carcinoma parenchyma) and spindle cells (79.2%) that had undergone an epithelial-mesenchymal transition(*41, 42*).

### Scaling gene networks from bulk tissue to single cells: metagenes quantify extended gene function

To visualize how metagenes are expressed at the single-cell level, we first examined *Lgr6* and its closely related family member *Lgr5*(*18*). Both genes are known stem-cell markers in the skin(*43, 44*) but *Lgr6* is clonogenic in squamous cell carcinomas whereas *Lgr5* is not. *Lgr6* as an individual gene was primarily expressed in the isthmus region of the upper hair follicle (arrow, Fig. 2a), and sparsely in the interfollicular epidermis. Prior lineage tracing studies have shown that *Lgr6*+ cells repopulate these regions during normal homeostasis and wound healing(*44, 45*). These cells then differentiate into the overlaying specialized epidermal strata. The *Lgr6* normal-skin metagene (the aggregate expression of the 100 genes most highly rank-correlated with *Lgr6* in bulk-tissue samples of normal skin) reflects this developmental pattern: increasing expression across interfollicular cells that peaks in terminally keratinized cells from the granular layer of the epidermis (arrow, Fig.2b). Furthermore, the individual genes that constitute the *Lgr6* normal-skin metagene are enriched for keratinocyte proliferation and differentiation (Data S5), implying that this network captures *Lgr6’s* function to establish epidermal cell layer patterning more strongly than its complex functions in other skin compartments such as the sebaceous gland or hair follicle(*46*). The *Lgr6* carcinoma metagene, on the other hand, shows high expression in the carcinoma parenchyma (arrow, Fig. 2c), reflecting the functional switch during tumor progression from normal homeostasis to malignancy (see also table S1). *Lgr5* is highly expressed in the lower bulge region of the hair follicle, reflecting its known follicular stem-cell function (arrow, Fig. 2d). Unlike *Lgr6*, the *Lgr5* carcinoma metagene shows low expression across all stages of carcinogenesis (Fig. 2e, fig. S2). Thus, metagene expression of the *Lgr6* carcinoma network increases, while that of the *Lgr5* carcinoma network decreases, in parenchymal cells during tumor progression (Fig. 2f,g). This therefore demonstrates that metagene expression in single cells mirrors the functional role of these stem-cell markers in normal tissue, and reflects functional rewiring that occurs during malignant transformation.

**Fig. 2.**
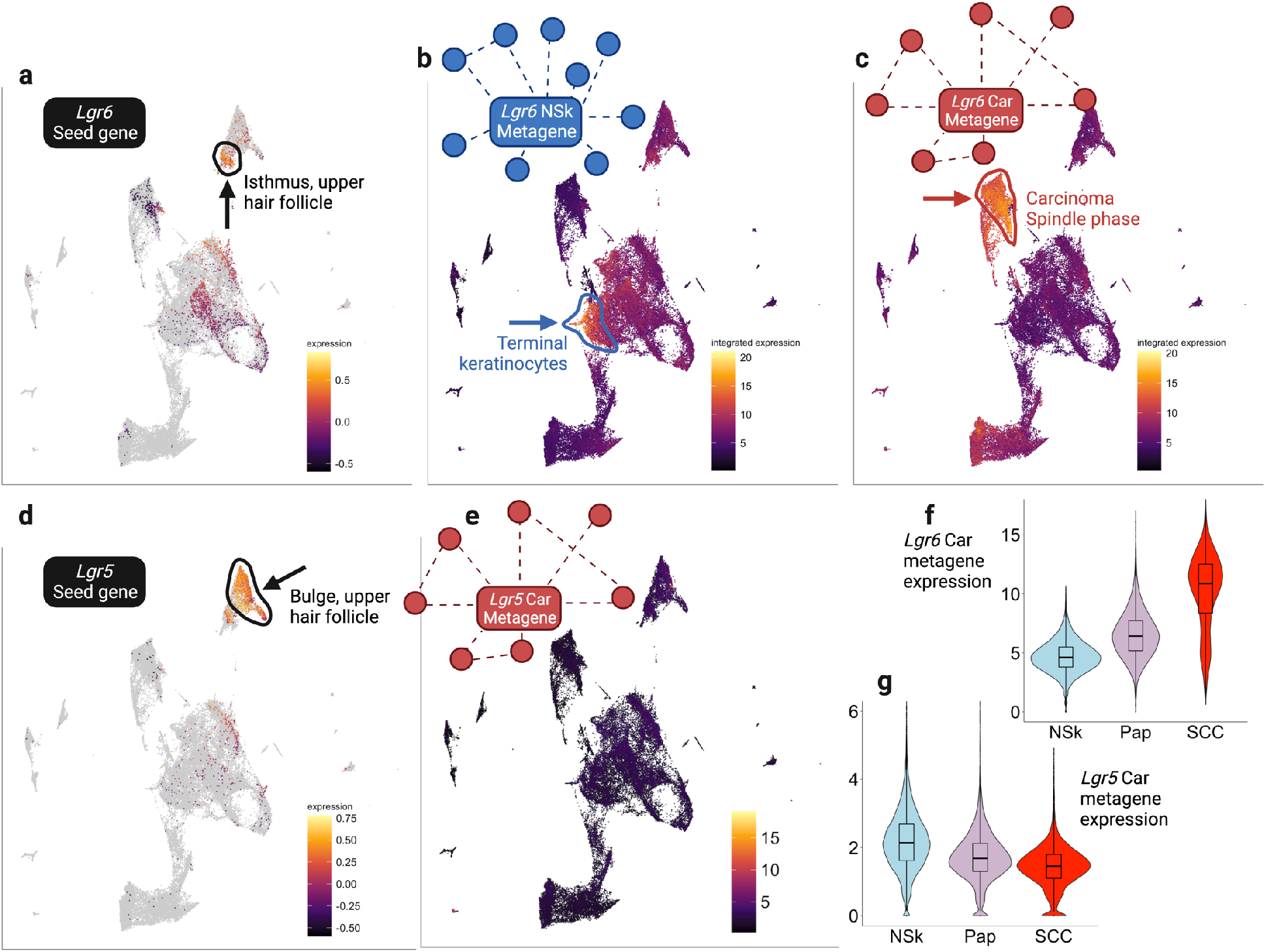
Single-cell expression of stem-cell seed genes and metagenes. (**a**) Expression of *Lgr6* alone in the single-cell UMAP. (**b**) Expression of the *Lgr6* normal skin metagene defined as the top 100-correlated genes to *Lgr6* in normal skin (NSk) bulk-tissue samples. (**c**) Expression of the *Lgr6* carcinoma metagene defined as the top 100 correlated genes to *Lgr6* in carcinoma (Car) bulk-tissue samples. (**d**) Expression of *Lgr5* alone in the single-cell manifold. (**e**) Expression of the *Lgr5* carcinoma metagene defined as the top-100 correlated genes to *Lgr5* in Car bulk-tissue samples. (**f**,**g**) A comparison of metagene expression during carcinogenesis for the *Lgr5* and *Lgr6* carcinoma metagenes. Y-axis is log metagene expression for each stage-specific parenchyma cell; box-and-whisker plots are median+IQR with outliers (Q1-1.5×IQR or Q3+1.5×IQR) not shown. All within-plot pairwise comparisons are significant at FDR<0.05.

### Convergent expression of multiple stem-cell metagenes in the same cell populations

Since Spearman groups 1 and 3 were strongly anticorrelated in tumor samples (Fig. 1b), we reasoned that they may represent two mutually exclusive stem-cell states. To test this hypothesis, we examined how stem cell markers, and their metagenes, were expressed in single-cell data.

The expression of individual stem genes was notably variable across single cells with sparse and divergent expression in disparate cell populations (seed gene panels in fig. S3). However, metagene expression dramatically altered this landscape. Spearman group 3 carcinoma metagenes co-localized to a specific “spike” cell population corresponding to unbiased cell cluster 33 in the UMAP of the papilloma (lower spike, Fig. 3). “Spike” refers to the appearance of this cell population in the UMAP. For example, carcinoma metagenes for *Pitx1, Krt15* and *Psca* were highly and specifically expressed in this lower spike population. In contrast, metagenes for *Bmi1* (Spearman group 1) or *Lgr6* (Spearman group 2) were almost absent from the lower-spike cell population (fig. S3). Although the upper spike showed higher expression for Spearman group 1 metagenes, this cell population showed distinct enrichment in expression of metagenes corresponding to markers of cell cycle progression, including *E2f1* and *Foxm1*, known as a master regulator of proliferation and malignant progression (Fig. 3) (*47*). Similar patterns were seen for multiple markers of DNA damage/genomic instability, for example those corresponding to *Atm* and *Atr* (Figs. 3 and S4). Together, we take this to indicate that the upper spike cells express high levels of gene networks related to mitosis and cell division.

**Fig. 3.**
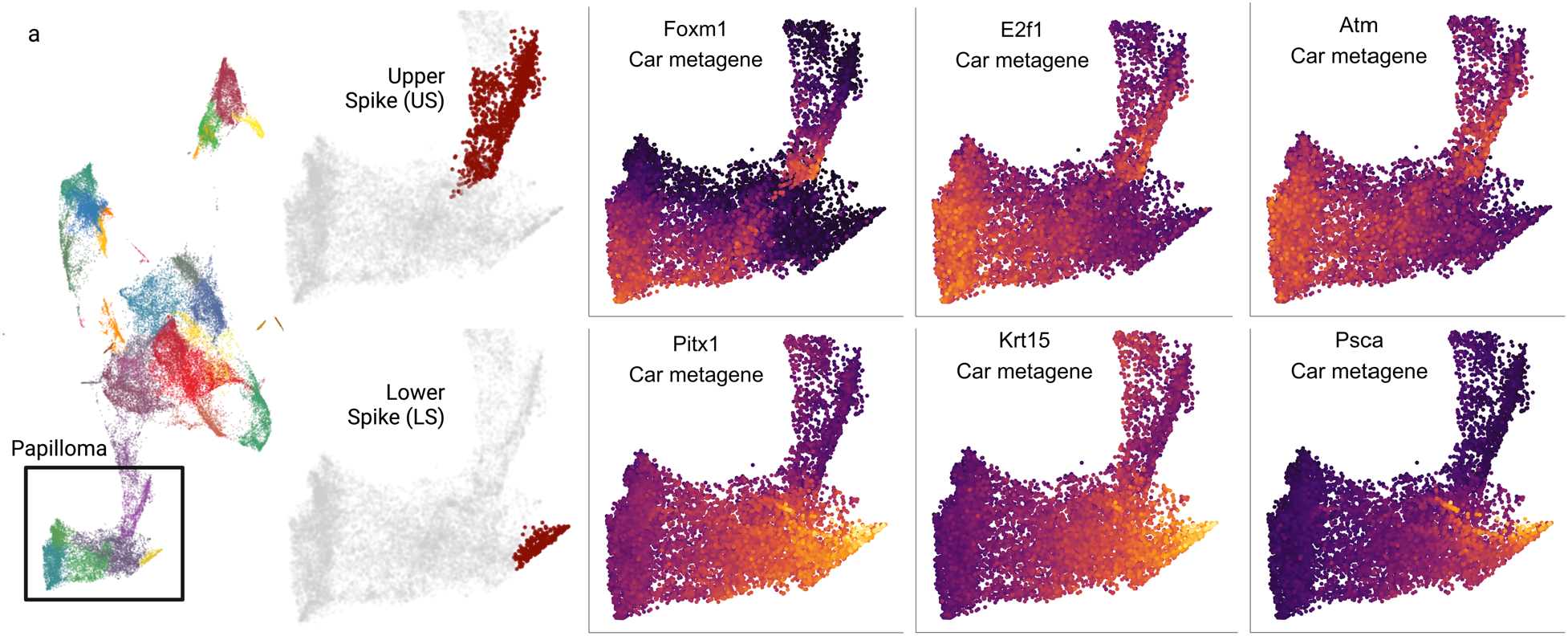
Exemplar carcinoma metagenes that localize to the upper spike (US, upper panels) and lower spike (LS, lower panels) of the papilloma. The location of the spikes is colored red in the panels next to the metagenes, and corresponds to unsupervised cell clusters 19 and 33. Purple is low expression and yellow is high expression.

We then hypothesized that if stem cell Spearman groups 1 and 3, which were negatively correlated at the bulk-tissue level, were truly mutually exclusive gene programs in the single-cell data, then their anticorrelated genes would be expressed primarily in mutually exclusive cell populations. We refer to the group of most highly anticorrelated genes as the ‘negative metagene.’ Strikingly, plotting Spearman group 1’s negative metagenes showed high expression in the lower spike region (fig. S5). This points to the mutually exclusive relationship between Spearman groups 1 and 3 carcinoma metagenes in these distinct single-cell populations.

To investigate possible overlapping functions of stem networks in the lower spike, we tested for gene ontology (GO) enrichment in genes of the ten stem-cell Spearman group 3 metagenes that co-localized to the lower spike. GO analysis showed that this set of 223 genes was highly enriched in functions related to oxidative stress (*Duoxa1/2*), skin barrier formation (*Sprr1a, Sprr3*), wound healing (*Wnt4, Klf5*), cell migration (*Cd44, Ceacam1*), apoptosis (*Anxa1*), and immune responses (*Arg1*) (table S2). Notably, normal wound healing is associated with oxidative stress and lineage infidelity, the latter indicative of stem cell plasticity and characterized by *de novo* co-expression of genes representing different hair follicle and epidermal lineages(*16*). Several genes associated with stem-cell plasticity were prominent in this set of lower spike metagenes (eg *Sox15, Foxa1, Klf5, Cd44, Wnt4*), consistent with the development of lineage infidelity in the lower spike region. Despite the individual seed genes being expressed in disparate cell populations, these well-known stem-cell markers converged at the metagene level with high expression in the same single-cell population, and this convergence cohered around lineage plasticity and wound healing.

### Alternative stem cell populations arise from *Lgr6*-positive papilloma cells

Having identified two distinct single-cell populations expressing high levels of alternate stem programs (the upper and lower spikes), we then asked how they relate to *Lgr6*, which is a major driver of clonogenicity in this system. We tested this directly through lineage tracing and immunofluorescence analysis of normal skin, papillomas and carcinomas (Fig. 4). In normal skin *Lgr6*+ cells repopulate the upper hair follicle and interfollicular epidermis (Fig. 4a), but during neoplastic progression, expression is more widespread and disorganized as *Lgr6* marks clone-initiating CSCs(*18*). In papillomas, rare *Lgr6-GFP+* cells were primarily located at the basement membrane, and their tdTomato+ progeny formed streaks that extended into the upper differentiated cell layers (Fig.4b)(*48*).Conversely, carcinomas showed highly disorganized tissue-level architecture (Fig. 4c).

**Fig. 4.**
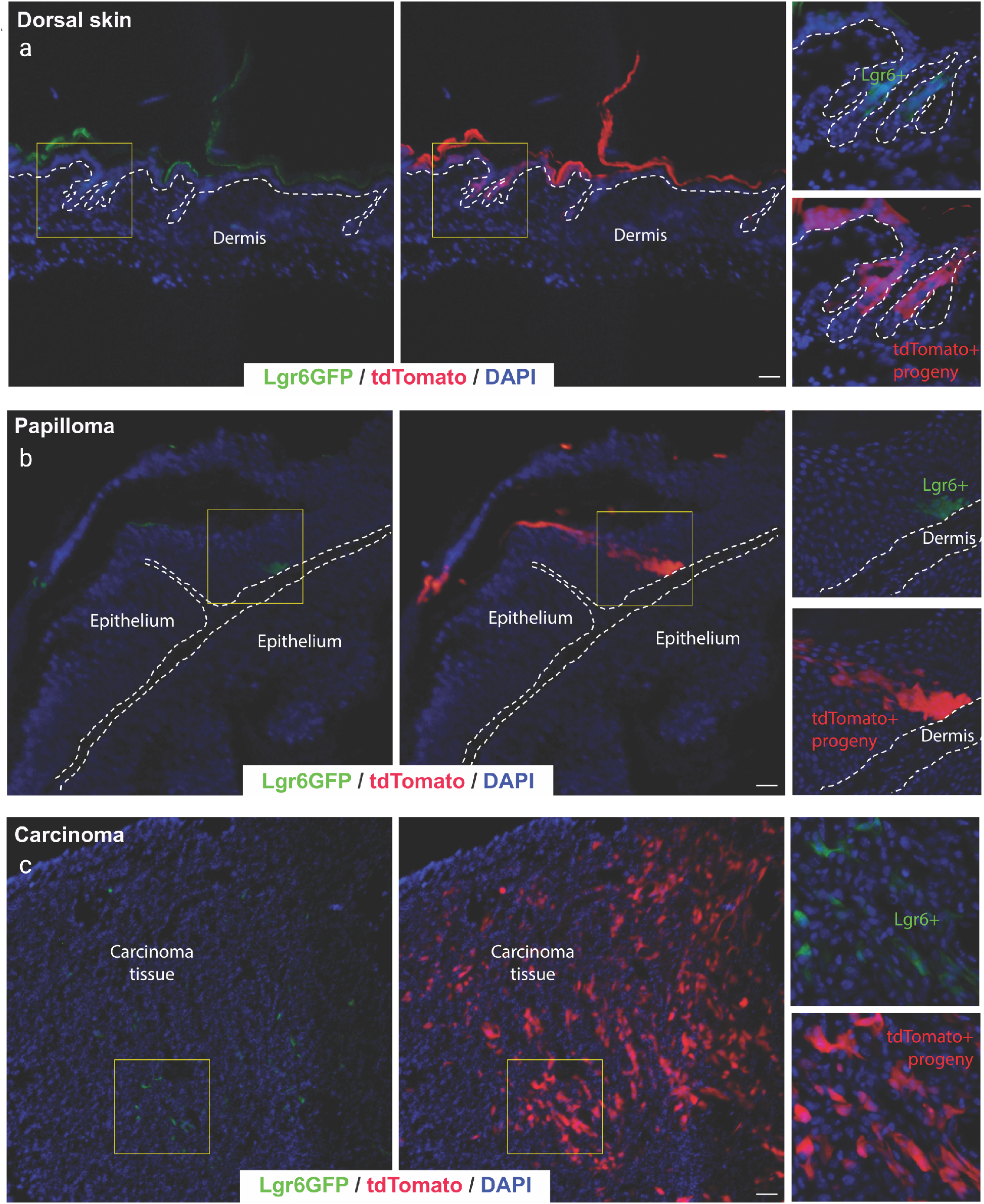
Immunofluorescent single-cell lineage tracing of *Lgr6*+ cells and their progeny. (**a**) Lgr6 Lineage tracing in normal dorsal mouse skin, benign papilloma, and carcinoma tissue. Representative immunofluorescence images of Lgr6-driven lineage tracing for: (**a)** normal dorsal mouse skin; (**b**) benign papilloma tissue; (**c**) carcinoma; 10 days after topical 4-OH-Tamoxifen treatment in vivo. *Lgr6*+ stem cells (green) are localized within (a), the hair follicle epithelium and epidermis of the normal dorsal skin; (b), predominantly the basal epithelium of the papilloma; (c), scattered throughout the carcinoma epithelium. These give rise to tdTomato+ (red) progeny within those tissues. Yellow boxes indicate the magnified region, dotted lines represent the epidermal-dermal border, and nuclei were counterstained with DAPI (blue), scale bar = 50um.

We then physically separated the *Lgr6*:GFP+ stem cells in normal skin, and in benign and malignant tumors, from their respective tdTomato+ progeny cells by flow cytometry, and examined the separate trajectories of these cells within the unsupervised cell clusters identified by scRNASeq (Fig. 5, fig. S6). Seven high-resolution clusters were identified in the papillomas, and clusters 19 and 33 corresponded to the upper and lower spikes, respectively (Fig. 5a). By examining the proportion of *Lgr6*:GFP+ cells and TdTomato+ progeny cells in each cluster, we observed that moving from cell cluster 14 to cluster 5 revealed a steady decrease in the relative number of *Lgr6*:GFP+ cells compared to their progeny cells (Fig 5a). From cluster 5 the spikes emerged with highly divergent *Lgr6* loads: the upper spike (cluster 19) was highly enriched for *Lgr6*:GFP+ cells and depleted in progeny cells, whereas the lower spike (cluster 33) showed the opposite pattern and was strongly depleted in *Lgr6*:GFP+ cells. In conjunction with the metagene patterns, these lineage-tracing results point to two highly divergent cellular fates: 1) the upper spike maintains *Lgr6*-associated self-renewal and proliferation, whereas the lower spike shows increased lineage infidelity, plasticity and commitment to different cell fates.

**Fig. 5.**
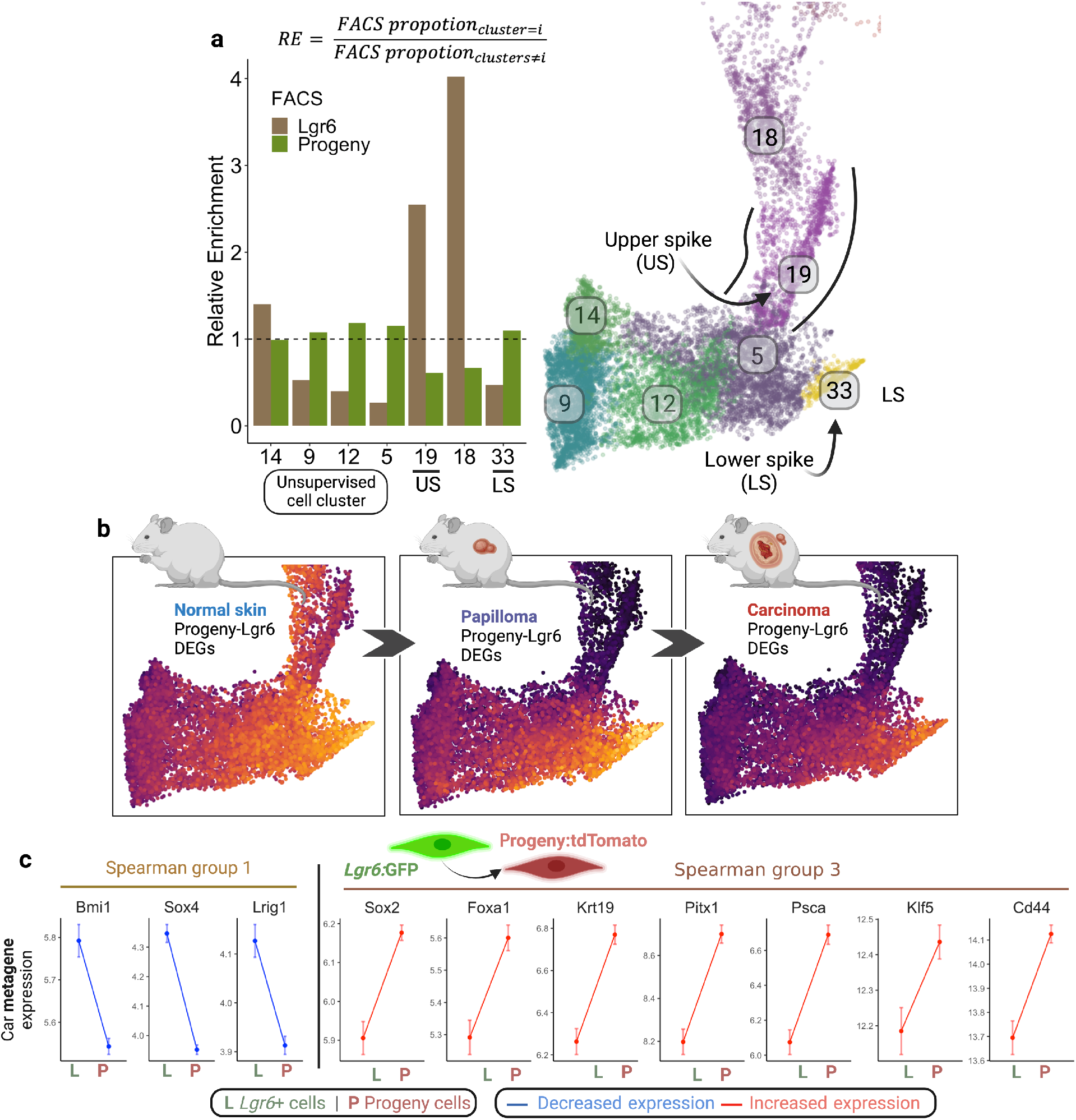
Lgr6-progeny cells form the lower spike and express high levels of a Spearman group 3 program. (**a**) Relative enrichment (RE) of unsupervised cell clusters for *Lgr6*:GFP or progeny:tdTomato cells sorted by FACS. Relative enrichment was calculated as the proportion of a cell type (*Lgr6*:GFP+ or progeny:tdTomato+) within the cluster divided by the proportion across all other cell clusters in the papilloma. For example, the relative enrichment of *Lgr6*:GFP+ cells in cluster 14 was the proportion of these cells in cluster 14 divided by the proportion in clusters 9,12,5,19,18, and 33. The UMAP of the papilloma shows high-resolution cell clusters with cluster numbers being arbitrary. (**b**) Aggregated expression of top 100 DEGs by fold change from progeny tdTomato+/ *Lgr6*:GFP+ cells discovered by testing for DEGs between normal skin parenchyma; papilloma parenchyma; and finally carcinoma parenchyma. (**c**) Carcinoma metagene (top) expression means across individual cells of the *Lgr6*:GFP+ (L) and tdTomato+ Progeny (P) lineages taken from neoplastic parenchyma cells (pooled papilloma and carcinoma parenchyma); red and blue lines are significant differences at Bonferroni-adjusted p<0.01 by t-tests, and grey are non-significant; error bars are standard errors.

We next performed a stage-specific differentially expressed gene (DEG) analysis comparing *Lgr6*+ and progeny cells (Data S6). At each stage (NSk, Pap, and Car), the DEGs that increased most in the progeny cells relative to *Lgr6*+ cells were most highly expressed in the lower spike (Fig. 5b). The overall intensity of expression however decreased from normal skin to papilloma, and then to carcinoma, in line with the known propensity for loss of differentiation capacity during malignant progression. We then tested how expression levels of specific stem cell metagenes change when *Lgr6*+ cells give rise to their progeny in benign or malignant tumors. Specifically, 7 lower spike-specific metagenes increased in expression from the *Lgr6*+ cells to progeny in tumor cells (Fig. 5c). Conversely, the 3 Spearman group 1 metagenes (*Bmi1, Sox4, Lrig1*) showed decreased expression across this trajectory. Taken together, these differential shifts in stem network expression across this empirically traced *Lgr6*→progeny lineage is strongly suggestive of a hierarchical relationship between stem-cell genes in these populations.

### Transition to the upper spike represents a cell cycle checkpoint

Our combined data from this and previous studies lead to a hierarchical model for malignant progression during multistage skin carcinogenesis. Lineage tracing studies have shown that papillomas become carcinomas by clonal expansion: single papilloma cells undergo a transition to increased growth and invasion (*48*). However, we demonstrated above that this transition is not universal at the single-cell level since *Lgr6*+ cells generate progeny that undergo one of two alternative cell fates: cells in the upper spike maintain *Lgr6*+ status, suggesting a self-renewal program, while the lower spike which is depauperate in *Lgr6+* cells undergoes growth arrest and differentiates along a number of different lineages controlled by a suite of different stem cell genes and lineage-specific transcription factors.

The observation that Spearman group 3 metagenes expressed in the lower spike cells are enriched in expression of markers of lineage infidelity, oxidative stress, apoptosis, and immune cell activation (table S2) suggested that these stress conditions could cause activation of tumor suppressor pathways. To interrogate this model, we examined the expression of metagenes for several well-known tumor suppressors, including p21/*Cdkn1a*, p16/*Cdkn2a*, and p15/*Cdkn2b* which are known to play important roles in keratinocyte growth, quiescence, and senescence in response to *Ras* activation(*49, 50*). All of these showed increased expression both at the single-gene and metagene levels in the lower-spike cells (fig. S4). Loss of the *p16/Cdkn2a/p15/Cdkn2b* locus on mouse chromosome 4 is a common event associated with progression to undifferentiated carcinomas in this model (*41*), as well as in human cancers(*51*). We conclude that analysis of *Lgr6*+ stem cell fate during pre-neoplasia at the single cell level illustrates features of alternative cell fates that control the balance between proliferation and differentiation/cell death.

### A conserved cell state expressing markers of drug resistance in mouse and human tumors

Since stem-cell plasticity in tumors has been commonly associated with resistance to chemotherapy or targeted drugs(*52, 53*), we sought to determine whether the lower-spike cells expressed empirically derived markers of drug resistance identified from drug screens. We examined markers of drug resistance in several human tumor types including basal cell carcinomas (BCC), prostate adenocarcinoma (PAD) and lung adenocarcinoma (LUAD). BCCs from patients treated with vismodegib(*54*) showed elevated expression of resistance genes *Tacstd2, Ly6d*, and *Lypd3*. We generated carcinoma metagenes for these seed genes using our mouse bulk expression data and visualized their expression in our mouse single-cell clusters. These BCC drug resistance metagenes were most highly expressed in the lower spike cells in papilloma (fig. S7). A high-plasticity cell state was also identified in human and mouse lung cancers that contributed to cell heterogeneity, drug resistance, and poor patient prognosis(*55*). The most significant genes marking this cluster of lung cancer cells were *Slc4a11* and *Tigit*, the latter being well known as a marker of immune responses and a cancer drug target. We generated mouse skin carcinoma metagenes for *Slc4a11* and *Tigit* and found that they too were significantly and specifically expressed in the lower spike (fig. S7). Finally, *Foxa1* has been identified as a marker of drug resistance that can be mutated in human prostate cancers and contributes to lineage plasticity during prostate adenocarcinoma development(*56, 57*). Although sporadically expressed as a single gene, its carcinoma metagene specifically localizes to the same lower spike region as other human genes associated with drug resistance (fig. S7). Together, these metagenes show that treatment-response genes found across highly disparate human cancers are most highly expressed in the same pre-neoplastic mouse cell population.

To directly test whether lower-spike gene networks are indeed upregulated during drug treatment, animals bearing primary carcinomas were treated *in vivo* with the commonly used chemotherapeutic agent *cis*-platin(*58*) and profiled by scRNAseq. We tested *cis*-platin-vs-control carcinomas for DEGs, finding 2461 significantly upregulated and 1298 significantly downregulated genes (Fig. 6a, Data S7). Upregulated genes were significantly enriched for cellular responses to stress involved in drug resistance such as reduction-oxidation, mitotic exit, and anti-apoptotic genes (Data S8). Strikingly, plotting the expression of the upregulated genes from these independent separate tumors into the original UMAP showed that they were most highly expressed in the lower spike (Fig. 6b,c).

**Fig 6.**
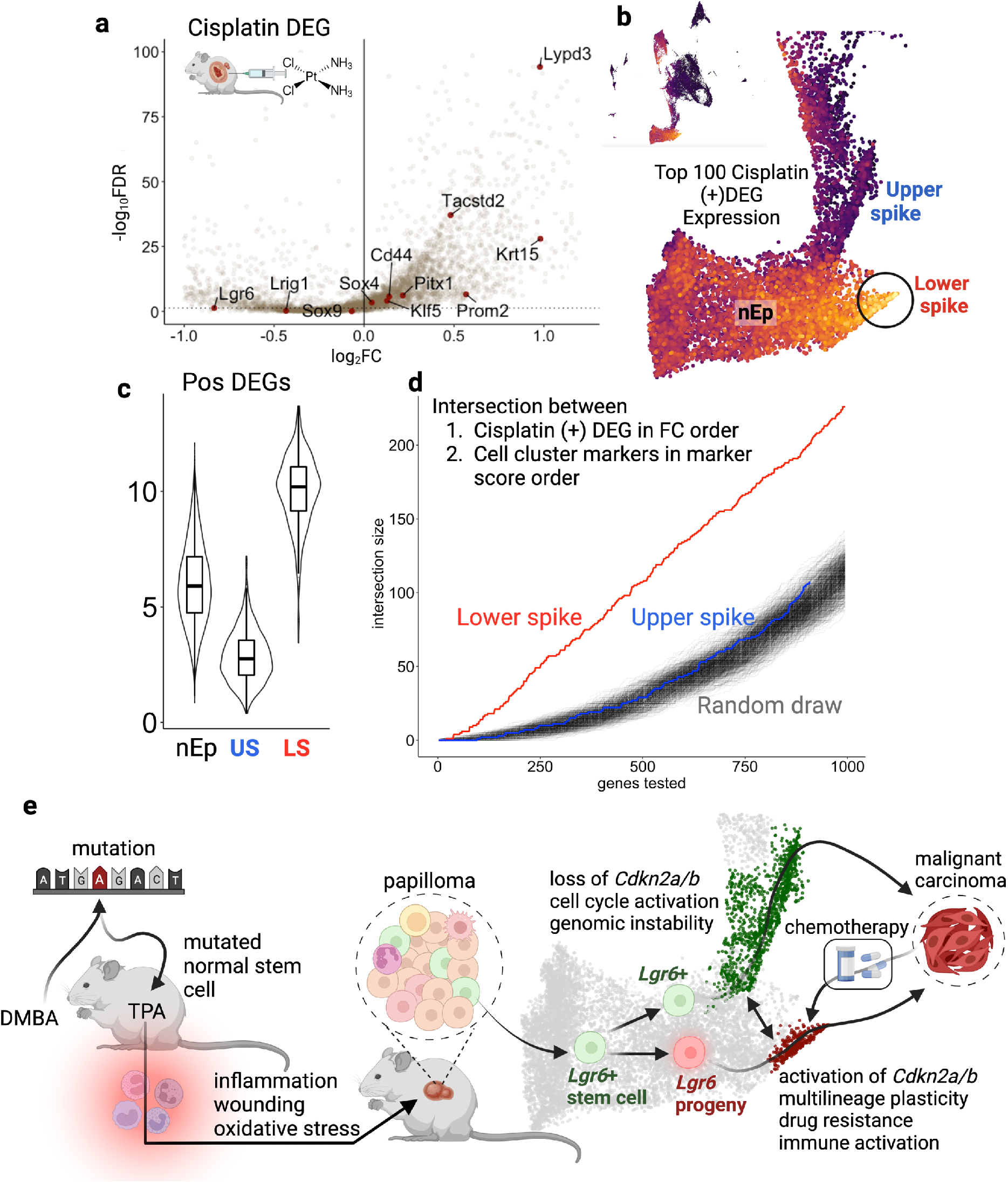
*Cis*-platin activates genes that localize to the lower spike. (**a**) Differentially expressed genes in squamous cell carcinoma parenchyma cells treated with *cis*-platin *in vivo*. Data derived from scRNAseq of 2 control and 2 treated tumors. Positive values represent genes that are upregulated by *cis*-platin, and negative downregulated. Labelled genes are those mentioned in the main text. (**b**) Integrated expression of the top 100 significantly upregulated genes according to fold-change. (**c**) Box and violin plots of integrated expression of the top 100 positively upregulated DEGs shown in panel b. nEp refers to all papilloma cells that are not in the spikes; US is for upper spike (cluster 19); and LS is for lower spike (cluster 33). All pairwise comparisons are significant at FDR<0.01. (**d**) The intersection of 1) genes that define a cell cluster ordered from high to low marker score and 2) genes that are upregulated by cis-platin challenge ordered from high to low fold change. The x-axis refers to the number of genes concurrently tested for intersection size in each list. For example, 100 refers to the top-100 marker genes in a cluster and the top 100 positively upregulated genes, which are then tested for the number of overlaps (intersection size) which is displayed on the y-axis. The red line shows the results for lower-spike markers; the blue for upper spike markers; and grey lines show randomly chosen sets of dummy marker genes. (**e**) The model for carcinoma development in single cells based on co-expression networks from bulk-tissue samples. DMBA initiates carcinogenesis by inducing oncogenic *Hras* mutations in cells which then lie dormant until promoted by the inflammatory promoter TPA. These show characteristic inflammatory and wound healing expression patterns as papillomas form. A single papilloma cell clonally expands and begins to undergo malignant transformation with a fraction of cells adopting a lower-spike cell state characterized by oxidative stress, reduced cell growth linked to elevated *Cdkn2a/b* expression, increased lineage plasticity, and immune activation; an alternative cell fate leads to the upper spike cell state characterized by loss of *Cdkn2a/b*, cell cycle activation, and genomic instability. Chemotherapy treatment of malignant carcinomas increases stress, causing a reversion towards the lower-spike high plasticity cell state.

Finally, we tested whether the overlap between DEGs upregulated by cis-platin treatment and lower-spike marker genes was greater than expected by chance. We used logistic regression to detect 1000 lower-spike markers, ordering them according to marker score which is a metric of the specificity of a gene’s expression for a particular cell population (Data S9). We then sorted *cis*-platin DEGs based on fold-change, and tested for the frequency of overlap between these two ordered lists. We established an empiric null by randomly drawing dummy marker sets of genes 1000 times and testing for overlap. Finally we repeated this analysis for upper spike markers.

Together this shows that the intersection between lower-spike markers and upregulated cis-platin DEGs exceeds random draws, but that the upper-spike intersection falls squarely within the null range (Fig. 6d). In other words, upregulated *cis*-platin DEGs are the same as lower-spike markers more often than expected by chance. Thus, both *cis*-platin DEG expression magnitude and overlap with lower-spike markers orthogonally validate our interpretation of the lower spike as a high-plasticity, chemotherapy-responsive cell population.

## DISCUSSION

Identifying and quantifying cell plasticity is an emerging challenge in cancer biology(*1, 2*). Numerous studies have identified a wide array of specific genes that contribute to plasticity signatures, but a common theme has emerged: at the level of individual cells, there appears to be a relatively small number of developmental paths towards malignancy(*55, 5, 60*). Here, we explored plasticity states *in vivo* in a direct and novel way by identifying expression networks for known stem-cell genes and tracing their expression across a continuum of stages of carcinogenesis at the single-cell level. The mouse model we used encompassed germline genetic heterogeneity, somatic genomic events and gene expression across multiple samples of normal, premalignant, and malignant tissue samples, to identify the steps necessary for rewiring of stem-cell networks at each stage. Importantly, tumors in this model were induced by sequential exposure to both mutagens and tumor promoters that elicit chronic inflammatory responses, factors increasingly seen to be critically important in human cancer etiology(*61*–*63*). We used this model to generate metagenes and analyzed their single-cell expression patterns, revealing clear developmental programs that are highly expressed in two distinct populations of pre-malignant cells associated either with quiescence and markers of stress-induced lineage plasticity, or with proliferation and self-renewal. Our data are reminiscent of bifurcations between proliferative and quiescent stem-like states that have been mechanistically traced *in vitro* to control of the cell cycle restriction point by balanced activities of *Cdk2* and the tumor suppressor gene *p21/Cdkn1a*(*64*).

Although plasticity in single cancer cells can be driven by stem-gene expression, there remains considerable controversy surrounding the relationships between stem-cell populations in normal tissues, and those that are found in tumors (*65*). One view is that a tumor stem-cell hierarchy exists (*66*), while others suggest that the chaotic, stressful environment associated with tumor growth induces extreme plasticity and heterogeneity, with no clear evidence of a hierarchical relationship between stem cells and their progenitors(*67*). Since decades of empiric effort have conclusively demonstrated that the stem-cell genes we have analyzed constitute *bona fide* stem-cell markers in skin, we reasoned that we could test how their relationships changed at both the bulk level through expression reorganization and at the single-cell level by measuring their expression in an empiric lineage-tracing system. At the bulk level, we uncover at least three distinct tumor stem-cell co-expression groups exemplified by 1) *Bmi1, Sox4*, and *Lrig1*, 2) *Lgr6* and *Sox9*; and 3) *Sox2, Pitx1, Klf5, Psca, Cd44, Prom1* and other well-known stem-cell markers. Spearman groups 1 and 3 appear to represent alternative states as their expression patterns are strongly anti-correlated, and as such may represent a major transition point in cells that undergo malignant conversion. In support of this possibility at the single-cell level, markers of Spearman group 3 are specifically and highly expressed in a subpopulation of pre-malignant cells characterized by lineage infidelity, oxidative stress, and wound healing. However, these cells also express high levels of counterbalancing gene programs such as protective immunogenicity, apoptosis, senescence, and tumor suppressor activity. The alternative cell state linked to Spearman group 1 (*Bmi1, Sox4, Lrig1*) lacks this protective antagonistic pleiotropy and is marked by increased expression of cell cycle and DNA damage response genes. Both cell states may arise from *Lgr6*+ stem cells through alternative self-renewal and lineage commitment cell fate decisions (Fig. 6e).

This does not imply that cells cannot travel in the opposite direction across this single step in the developmental hierarchy during carcinogenesis. In fact, it is increasingly clear that cancer stemness is not bound by the same rigid unidirectional hierarchy that exists in most normal tissue since cancer stemness responds to complex microenvironmental interactions(*68*). In fact, we demonstrate that a range of human tumor types, as well as malignant mouse skin primary tumors, respond to chemotherapy *in vivo* by returning to this conserved high-plasticity lower-spike state that is a major feature of untreated premalignant papillomas. Together, these constitute unexpectedly complementary and orthogonal findings that integrate disparate observations in the literature regarding cancer stem cells and drug resistance, and provide a method that can be used by the community to interrogate expression of dysregulated cancer gene networks, rather than just single genes, at the single-cell level.

## Acknowledgments

We are very appreciative of the helpful discussion and comments from our colleagues to help refine our study and thank the UCSF Laboratory for Cell Analysis (LCA) Shared Research Facility, supported through the NIH grant P30CA082103, for flow cytometry assistance. We thank Natasha Carli of the Gladstone Genomics Core for scRNAseq library preparation and quality control for sequencing and Eric Chow for his assistance with sequencing at the Chan Zuckerberg Biohub facility.

## Funding

National Cancer Institutes (NCI) grant R35CA210018.

NCI grant R50CA25147 (EK)

UCSF School of Medicine Deep Explore (MT)

US National Cancer Institute (NCI) RO1CA184510

NCI Grant UO1 CA176287

Barbara Bass Bakar Professorship of Cancer Genetics (AB)

Cancer Research UK Mutographs Cancer Grand Challenge Award (C98/A24032)

CRUK/NCI Prominent Cancer Grand Challenge Award

## Author contributions

E.K. R.J.A and A.B. conceived and designed experiments. E.K. performed lineage tracing, histological imaging, and scRNAseq preparation and quantification. M.T. performed quantitative analyses of scRNAseq data. K.H., R.D., D.W. and O.M. carried out mouse studies, genotyping and isolation of RNA and DNA. M.K., H.G., D.Q., Y.R.L. and S.B. provided bioinformatics and computational support. Data analysis was carried out by M.T., E.K., R.J.A and A.B. M.T., E.K. and A.B. wrote the manuscript with input from all authors. A.B. supervised the overall project.

## Data and materials availability

All code to analyze the data are available on GitHub at https://github.com/maktaylo/Metagene-paper.

## Supplementary Materials

Materials and Methods

Supplementary Text

Figs. S1 to S7

Tables S1 to S2

