## Supplementary Material for "Gene networks reveal stem-cell state convergence during preneoplasia and progression to malignancy in multistage skin carcinogenesis"

Science
AAAS

### Supplementary Materials for

#### Gene networks reveal stem-cell state convergence during preneoplasia and progression to malignancy in multistage skin carcinogenesis

Mark A. Taylor, Eve Kandyba, Kyle Halliwill, Reyno Delrosario, Matvei Koroshkin, Hani Goodarzi, David Quigley, Yun Rose Li, Di Wu, Saumya Bollam, Olga Mirzoeva, Rosemary J. Akhurst, Allan Balmain

**The PDF file includes:**

Materials and Methods
Supplementary Text
Figs. S1 to S7
Tables S1 to S2
References

**Other Supplementary Materials for this manuscript include the following:**

Data S1 to S11

**Materials and Methods**

No power analysis was used to pre-estimate sample sizes prior to analysis. Experimental mice were randomized for DMBA/TPA treatment. Investigators were not blinded to mouse treatment factor levels during analysis.

Mouse husbandry

Animals were housed under standard conditions, fed *ad libitum* and treated in accordance with the rules and protocols stipulated by the UCSF Institutional Animal Care and Use Committee (IACUC). All mouse experiments were approved by the University of California at San Francisco Laboratory Animal Resource Center.

Chemical Carcinogenesis experiments

For chemical carcinogenesis experiments, 8 – 12 week old male and female mice were randomly assigned to control or treatment groups. Initiation was carried out using a single dose of the mutagen dimethylbenzanthracene (DMBA, Sigma, D3254, 25 ug per mouse in acetone) applied topically to shaved back skin. After 7 days, the mice were administered biweekly topical treatments of 12-O-tetra-decanoylphorbox-13-acetate (TPA, Sigma, P8139, 200 ul of a $10^{-4}$ M solution in acetone) for 20 weeks. The animals were then monitored for papilloma number and progression to carcinoma. Skin tumors were surgically resected in accordance with IACUC guidelines and mice were euthanized per animal requirement with dorsal skin, visible papillomas and carcinomas collected for analysis. Mouse and sample information are summarized in Supplementary Table 10.

Bulk RNA expression experiments

Male SPRET/Ei and female FVB/N mice were obtained from the Jackson Laboratory, female F1 hybrids were crossed with male FVB/N mice to produce a 720-member heterogenous F1 backcross cohort for chemical carcinogenesis experiments. For bulk expression analysis of carcinomas, the entire lesion was excised, surrounding normal skin was trimmed away and the sample was snap frozen and stored at -80ºC. Normal skin from independent mice under the same experimental conditions was also sampled. To extract mRNA, frozen tissue was ground by chilled mortar and pestle, and suspended in TRIzol. TRIzol RNA extraction was then performed, followed by purification by Qiagen kit. mRNA was assayed on Affymetrix M430 2.0 chips according to manufacturer protocols.

Tamoxifen-induced, *in vivo* fluorescent Lgr6 stem cell-derived lineage tracing

To permanently fluorescently label Lgr6+ skin stem cells and all daughter progeny for lineage tracing experiments in skin and tumors, two doses of 4-OH tamoxifen (Sigma, T5648, 25 mg/ml in 100% ethanol; 5 mg per dose) were administered to shaved *Lgr6eGFPtdTomato* mouse back skin on the same day (once in the morning and at the end of the day). When skin tumors were present, the same tamoxifen dosage was topically applied directly over the lesions and onto the surrounding tumor adjacent skin. Lineage traced skin and tumor samples were collected 10 days after the initial tamoxifen treatment.

Tissue collection and confirmation of tdTomato+ in dorsal mouse skin and skin tumors

To preserve *Lgr6*GFP and tdTomato fluorescence within lineage traced tissue, prior to embedding, a small piece of the excised skin / tumor used for analysis was fixed in 4% paraformaldehyde (Electron Microscopy Sciences, 15710) for 2 hours and then placed in 30% sucrose (EMD Millipore, 573113, w/v in PBS) overnight at 4°C before freezing in OCT (Tissue-Tek, 4583) and storage at -80C. Analysis of skin *Lgr6*GFP stem cell-derived lineage tracing was verified using cryosection analysis for the presence of tdTomato+ (red fluorescence). For cryosectioning, fixed OCT embedded samples were sectioned at 10um thickness onto SuperFrost slides (Fisher, 12-550-15) prior to antibody staining and imaging with a Zeiss Axiovision microscope.

Immunofluorescent staining

Slides with 10μm cryosections were brought to room temperature and washed in 1x PBS (Invitrogen, AM9625) containing 0.1% Triton X-100 (v/v in 1x PBS) for 30 mins. Sections were then blocked using 10% normal goat serum (Jackson ImmunoResearch, 005000121) in 0.1% triton X-100 / PBS for 30 mins at room temperature and then incubated with a primary anti-GFP antibody (Abcam, ab13970) for 3 hours at room temperature. Slides were then washed 3x in PBS containing 0.1% triton X-100, and then incubated with a fluorescent secondary antibody (Invitrogen, A11039) for 60 mins at room temperature in the dark. Sections were then washed 3x in PBS containing 0.1% Triton X-100, counterstained with DAPI solution (Invitrogen, D3571) and mounted using Fluoromount mounting medium (Sigma, F4680).

Preparation of single cell keratinocyte suspensions for flow cytometry

Murine dorsal skin tissue (with subcutaneous fat removed) and tumors from Lgr6eGFPtdTomato mice were processed immediately following procurement. Samples were processed from 8 week old, untreated dorsal back skin (3 pooled female back skins); 8 week old, tamoxifen treated dorsal back skin (with 10 day lineage tracing); DMBA/TPA treated back skin adjacent to skin tumors; benign papillomas (4 pooled tumors) and carcinoma tissue (from one single carcinoma, trimmed of visible normal skin) prior to flow cytometry. The adjacent skin, benign papillomas and carcinoma tissue were collected from a single male Lgr6eGFPtdTomato mouse (with 10 day lineage tracing). Single cell suspensions of dorsal back skin were prepared by carefully removing subcutaneous fat and floating the intact tissue dermal side down in 0.25% Trypsin/EDTA (Invitrogen, 25200-056) solution with gentle rocking for 1 hr at 37°C. Trypsin activity was neutralized with 10% chelexed FBS in Ca2+ free PBS (Hyclone, SH30028.02). The epithelial skin layer was then scraped into suspension and agitated to dissociate cells. To obtain single cell suspensions of skin tumors, lesions were finely chopped into very small pieces with a sterile scalpel and then enzymatically digested in 0.25% Trypsin-EDTA with gentle agitation for 1 hour at 37°C. Individually, each disaggregated cell suspension sample was then passed through a sterile 40μm filter, centrifuged at 300g for 5 mins to pellet cells and resuspended in 10% chelexed FBS in Ca2+ free PBS prior to flow cytometry. Cell samples were collected and analyzed using a FACSAria III, and appropriate gating was used to exclude doublets. DAPI staining was used to assess cell viability and only viable, DAPI negative, cells selected for subsequent analysis. Within each sample type three populations of viable cells were evaluated: unenriched global cell populations, *Lgr6*GFP+ stem cells (gated on positive GFP expression), and tdTomato+ *Lgr6*+ stem cell-derived progeny cells (gated on positive tdTomato expression). FACS sorted cells were collected in Ca2+ free, PBS containing 1% chelexed Ca2+ free FBS, centrifuged at 300g to pellet the cells before final resuspension in 35µl 1% chelexed Ca2+ free PBS and storage on ice prior to scRNA-sequencing library preparation which occurred within 2-3 hrs of sample collection.

<u>Cisplatin chemotherapy experiment</u>

For the single-cell analysis of carcinomas treated by *cis*-platin, we generated four carcinomas as described above ("Chemical Carcinogenesis experiments") in 3 independent female mice. Chemotherapy treatment began when visible carcinomas reached 1 cm in diameter and consisted of intraperitoneal injections. Two mice bearing primary carcinomas were treated with 100uL of 0.8mg/mL *cis*-diammineplatinum(II) dichloride (D3371, TGI Chemicals), and one control mouse with 2 primary carcinomas was treated with 100 uL of PBS. Treatments were carried out twice, separated by 4 days, for both control and *cis*-platin treated mice. Treated carcinomas were harvested 3 days after the second treatment, and scRNAseq libraries were prepared as below.

<u>Single-cell RNA sequencing (scRNAseq) library preparation</u>

scRNAseq libraries for samples were prepared using the 10x Genomics Chromium extraction and preparation protocol according to the manufacturer's instructions and sequenced on a Novaseq S4 sequencer (Illumina).

<u>Sequencing read alignment, assignment, and quality control</u>

Reads generated by 10x Genomics sequencing were demultiplexed using default settings by Cell Ranger's mkfastq function (version 2.0, 10x Genomics). Demultiplexed reads were aligned and assigned to the *mm10* (GRCm38) mouse reference provided by 10x Genomics and converted to sample-specific unique molecular identifier (UMI) count matrices using the count function with default settings. Count data for each sample were independently quality-controlled. Minimally, all cells were required to express at least 200 genes, with at least 20% of reads aligning, and retained genes were required to be detected in at least 3 cells. We filtered stressed or dying cells with more than 10-17% of UMIs generated from mitochondrial genes (see Data S10). Some tissues, such as rapidly cycling normal skin or cancer cells, are highly metabolically active and thus may have these relatively high mitochondrial transcript content without it being an indication of cell inviability. We then analyzed sequencing curves of cells ordinated by UMI detection and gene number in order to detect lower elbows indicating low-complexity libraries (empty droplets) and upper elbows indicating an artifactually rapid increase in well-wise UMI detection (multiplets). We filtered cells falling below and above these thresholds, respectively. The lower threshold ranged between 250-750 and the upper between 2500-6000 (Data S10).

Normalization, batch correction, clustering

We integrated and normalized count data using R/Seurat v.2.3. To normalize, we used the NormalizeData function to scale UMIs by dividing each cell by the total number of counts per cell, multiplying by a scaling factor of 10000, and converting to a log scale. We then passed these data to the R software package Monocle3. We then included all genes in dimensionality reduction with principal components, limiting the reduced hyperspace to 100 dimensions. Together, the first 3 PCs explained 34% of the data, so the data were subsequently projected onto a 3-D manifold using the uMAP method(*1*). However, upon inspection of sample arrangement in this space, we observed that FACS was strongly confounded by batch. In particular, unsorted cells were colinear with the first batch, and sorted cells with the second batch. To reduce this technical variability while preserving biological variability, we implemented a mutual nearest neighbour method to identify hyperdimensional batch-specific planes and resolve these planes into a single mutually uniform plane(*2*). We then used these batch-corrected data for all downstream analyses. To detect unsupervised clusters of cells on different scales of transcriptomic similarity, we implemented the Leiden community detection algorithm within Monocole3 to call low-resolution clusters (referred to as "partitions" by the monocle software suite) and high-resolution clusters (referred to as "clusters")(*3*).

Cell type annotation

First, we implemented an unsupervised, bottom-up approach to identify genes that are highly and specifically expressed by high-resolution cell clusters. To do this, we used R/monocole3/top_markers to implement logistic regression wherein each high-resolution cluster was a predictor, and examined the top 50 most specific genes for each cluster, manually reviewing these sets for cell type. Next, we compiled a custom database of dermal, epidermal, stromal, and iimmune marker genes based on a literature review of publicly available mouse markers sets (Data S11). With this database, we trained a hierarchical classifier based on cross-validating 100-n random partitions of the entire data, and then applied the trained classifier to the entire dataset simultaneously(*4*). Second, we used this database to construct cell-type gene signatures and examined the relative expression of these signatures in the manifold. Together, these methods consensus-classified 57571 (99.591%) cells. We carried out high-resolution cell type deconvolution for 1) the carcinoma parenchyma identifying its squamous and spindle phases separated in its own manifold; 2) macrophage-Langerhans cells to test for the presence of M1-M2 polarization and transition among these cell types (Extended Data Fig. 1d); and among T cells to test for Cd4/8 differentiation (Extended Data Fig. 1e).

WGCNA gene clusters, Spearman gene clusters, metagenes

We implemented WGCNA on our normal skin and carcinoma samples separately. For each, we tested soft-thresholding powers between 1 and 20, choosing 6 for both. Minimum module size was 30 and max was 4000, with default clade reassignment and cut height parameters. For our 16 seed genes of interest, we calculated the Spearman rank correlations between these genes and all others present in the transcriptome. Since the correlations were significantly stronger in Car than NSk ($\overline{\rho}$=0.21 and 0.12, respectively [p=0.016, 1-sided t-test]), we searched for clustering patterns among seed genes in the Car correlation matrix using R/hclust with the complete method. This resulted in seed clusters 1-3, with the exception of Sox2 which was originally grouped in cluster 2 but which we considered to be in cluster 3 due to its similar single-cell metagene pattern as explained in the text. Next, we inferred rank correlation co-expression networks from bulk data, terming their aggregate behaviour as metagenes in the single-cell data. To do this, a seed gene was chosen and Spearman correlation coefficients to all genes in the microarray were calculated. The top 100 correlated genes were taken as the positive network, and the top 100 anticorrelated genes as the negative network. Next, we identified those genes expressed in the single-cell data, log-normalized their expression values with a pseudocount of 1, and scaled and summed them. This produced a single integrated value representing the aggregate expression level of the 100n network, and we considered this to be the metagene expression value.

Seed and metagene visualization and comparison

We overlaid $\log_2$ transformed individual gene and metagene expression on individual cells on dimensions 1 and 2 of the 3D manifold. In order to simultaneously compare expression of metagenes among specific cell groups and among metagenes as in Fig2b, we first divided the parenchyma into 7 tissue-specific cell groups shows in Fig.2a; next we aggregated gene expression for all genes within those cell groups using R/monocole3/aggregate_gene_expression; we then summed group-aggregated expression values across all genes within an metagene, dividing this value by the number of constituent genes; finally, we centred and scaled each row (metagene) independently. For consistency with the seed clusters, row order was maintained as in the Car seed gene cluster (Fig1b). To compare seed and metagene expression across the Lgr6→progeny hierarchy, we isolated stage-specific parenchyma cells; calculated metagene expression within each cell; tested for differences in the distribution of metagene and seed gene expression values with t-tests, adjusting significance for multiple testing with Bonferroni corrections (n=14).

GO Analysis

We implemented R/topGO v2.44.0(*5*) to test for functional enrichment in gene sets using Fisher exact tests, correcting for multiple testing with Benjamini-Hochberg, and comparing to genome-wide annotations provided by R/org.Mm.eg.db v3.8.2(*6*). We also tested for enrichment of genes sets using multiple gene annotation compendia compiled by the STRING database(*7*), including KEGG, Reactome, UniProt, Pfam, SMART, and InterPro, and tested for enrichment using the R package STRINGdb (*8*). Since resulting gene annotations were largely similar, we report FDR significance in enriched terms taken from the STRING GO categories.

Chemotherapy scRNAseq analysis

We identified differentially expressed genes (DEGs) between control and cis-platin carcinomas by first isolating carcinoma parenchyma as above. We then narrowed candidate genes to those expressed in at least 10% of parenchyma cells, and among those, to the top 5000 most variable genes using Seurat's FindVariableFeatures with the vst selection method. Next we used non-parametric Wilcox rank sum tests to compare expression between the control and cis-platin carcinomas, . In order to test whether DEGs from the cisplatin experiment were over-represented in markers genes for the spikes, we first used Monocole3's top_markers function to identify markers for the lower spike (unsupervised high-resolution cell cluster 33) and the upper spike (unsupervised high-resolution cell cluster 19), comparing these against all other cells present in the experiment and testing one thousand genes per group. We then ordered spike-specific marker genes from high to lower marker scores, and ordered significant DEGs from high to low fold change. We then scanned each list for intersection between at simultaneous rank, for example asking how many overlaps exist at the top 10 for each, then top 11, etc. In order to establish an empiric null distribution of overlaps that would be expected by chance, we repeated this procedure 1000 times with 1000 dummy marker sets consisting of randomly chosen 'marker' genes.

#### Supplementary Text: Defining Network Metagenes

We present here the rationale for our definition of network metagenes; describe methods used to determine an informative network metagene size (i.e. the number of genes included within a network metagene); and examine metagene expression stability across sizes.

**Transcriptomic landscapes are simplified by gene networks**. Relationships among genes are often quantified as networks, sometimes called co-expression or gene regulatory networks(*9*). The concepts, language, and methods to infer gene networks are diverse and often conflicting(*10*). At their core, however, is a common premise that an individual gene is embedded within a functional module and that it correlates to other genes within this module, which together form a network(*11*). The identification of such networks exploded in popularity due to easily implemented methods that factor co-expressed genes into their most-informative low-dimensional vectors (eigenvectors)(*12*), referred to as metagenes(*13*). We adopt this terminology, referring to our gene networks when we discuss their behavior as a single aggregate entity as network metagenes. Such networks have been extensively explored in bulk-tissue transcriptomes, and have been shown to control cell fate decisions more powerfully than individual genes(*14*). Furthermore, stem network dysregulation during oncogenesis has been convincingly demonstrated in bulk-tissue assays(*15*). Notably, until recently, metagene analyses have exclusively been carried out in bulk-tissue datasets, often comprising hundreds to thousands of samples. These robust sample sizes allow for high confidence in any particular metagene and its networks dynamics. However, bulk-tissue transcriptomics obscures gene expression which occurs within individual cells since it averages over all of the cells contained in the bulk sample. This reduces power to discern transcriptional patterns in specific cell populations such as stem cells which constitute only a fraction of any tissue(*16*).

**Scaling gene networks from bulk tissue to single cells.** Methods to infer gene networks from within single-cell data are often limited to *a priori*-derived transcription factor sets(*17*), and are especially sensitive to technical error and remarkably poor replicability across statistical frameworks, tissues, and samples within the same tissue or study(*18–21*). However, the weakness of gene-gene correlations in single-cell datasets(*18*) can complemented by the much stronger correlations in gene networks uncovered by bulk data(*22*, *23*). We therefore reasoned that we could complement our scRNAseq data with our much larger bulk expression database. To do this we leveraged our bulk tissue biobank to generate co-expression networks that were downscaled to the single-cell level. Our goal was *not* to define, assay, or infer the most important or parsimonious gene network in any single-cell population. As discussed above, exercises in network inference and empiric testing are open to a number of conceptual and methodologic concerns. For example, gene networks change across different cell populations—which cell population should a network be expressed in to be considered an important functional driver? Which modules within a functional network should be considered, and how should these considerations be constrained (ie functional constraints, correlation threshold constraints, mutual information constraints, etc)? The purpose of this study is not to provide answers to these questions since, although important, they warrant an extensive program of experimentation, analysis, and interpretation. Instead, we use metagenes to sample networks that are operative in previously unknown ways in single cells to gain novel insight into multi-gene expression dynamics during carcinogenesis in this system.

**Metagenes encompass previously ascertained networks**. A major question is to what degree our definition of metagenes captures functionally relevant networks. Since gene function is carried out at the protein level, we sought to understand how metagenes reflected both gene-gene and protein-protein networks. Similarly to gene expression-based network, construct validity in assaying and inferring protein networks is multifarious and often sharply conflicting(*24*, *25*). In order to remain agnostic to this debate, we used v11 of the STRING platform(*7*) to tune our metagene definitions. This platform aims to “collect, score, and integrate all publicly available sources of protein-protein interactions” and encompasses co-expression, text-mining, biochemical/genetic data, and previously compiled protein interaction databases. We first asked how connectivity (number of total edges) changed as a function of node number. To do this, we expanded metagene sizes for each of our seed genes from the top 5 to the top 500 most correlated genes in increments of 5, and assessed the total number of unique all-vs-all connections with a STRING score of at least 200. We then plotted this on a semi-log scale (Fig.SM1), which revealed that connectivity relates node size according to a power law for each seed gene. Although the shape of this power law differed among seed genes, we observed a remarkably consistent inflection point near node size N=100 in the growth of new connections. This was a first indication that N=100 may be an appropriate metagene size.

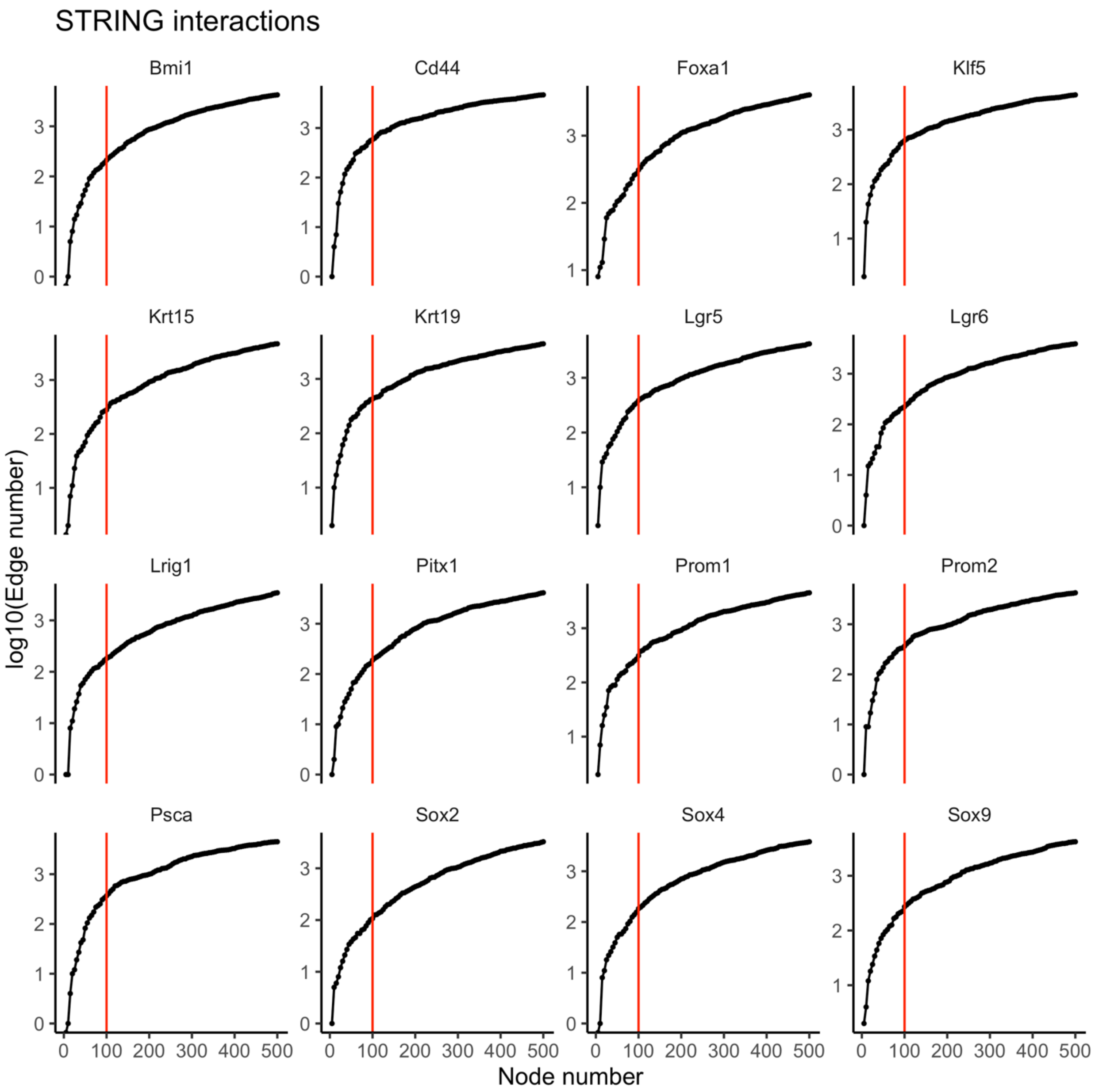


**Fig. SM1 |** Node number versus STRING edges. Panels are for separate tumor metagenes, seeded by the gene shown in the panel title; the x-axis is the number of genes included within an metagene in increments of 5 from the top 5 to top 500 most correlated genes to the seed gene; y-axis shows the $\log_{10}$ number of protein-protein network associations indexed by STRING among all of the genes included in an metagene of the size denoted by the x-axis; red vertical lines are for N=100 metagenes.

We then used the STRING database to assess whether metagenes at N=100 encompassed more network connections than would be expected by chance. For example, for *Lgr6*, 218 previously ascertained interactions were detected among the 97 gene models that mapped to those genes (Fig.SM2). The null expected number of interactions for this 97-member protein set was 113. We then tested whether this was a significant enrichment using the function get_ppi_enrichment in the R package STRINGdb v2.4.1(*8*). We repeated this for the 16 selected stem and cancer driver genes shown in Fig.1b, and all metagenes were significantly enriched for network interactions (Table SM1).

proteins: 97
interactions: 218
expected interactions: 113 (p–value: 0)

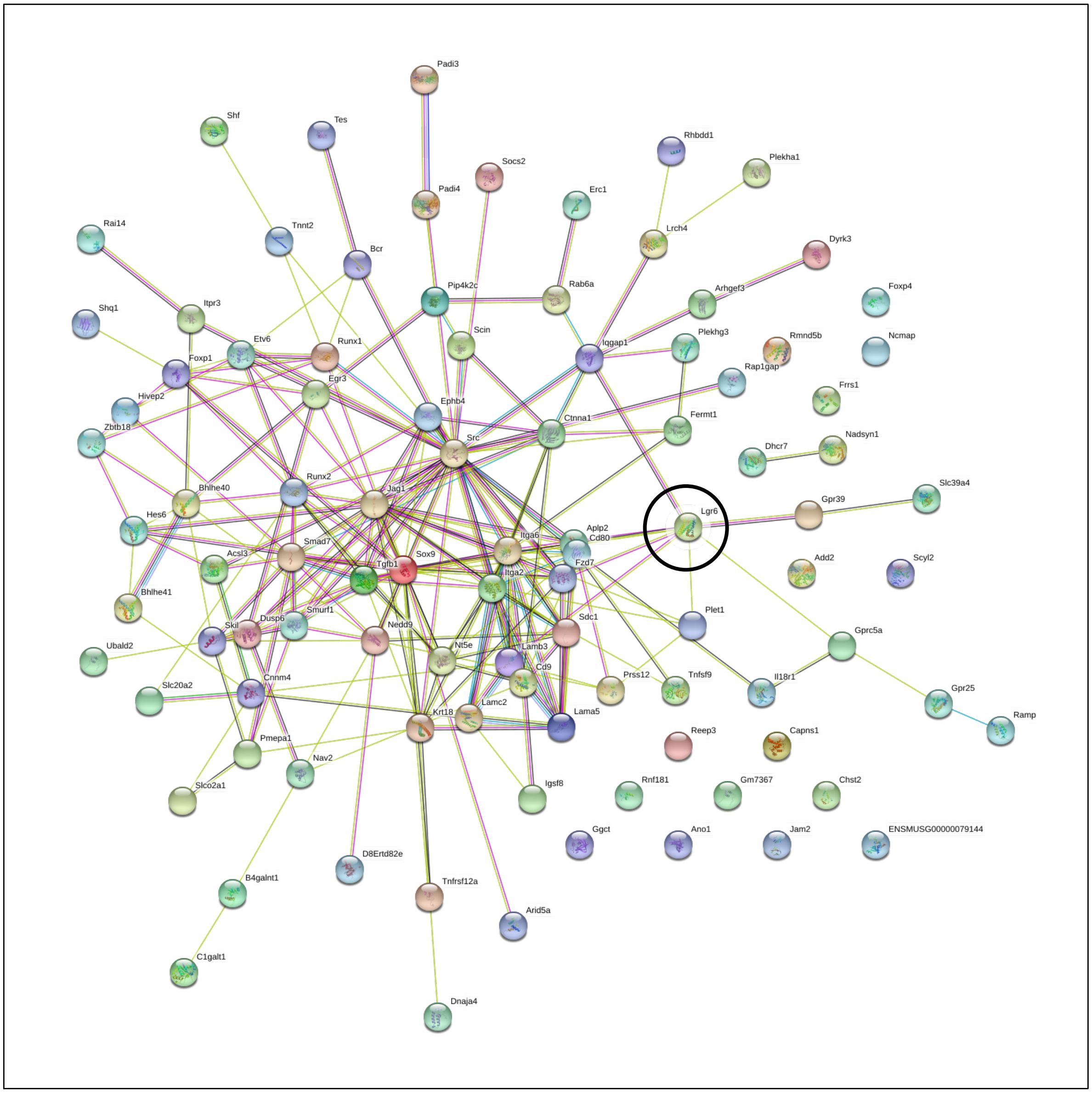


**Fig. SM2 |** Interactions (edges) among proteins encoded by the 97 genes mapping into the STRING database of the carcinoma metagene seeded by *Lgr6*. The black circle shows the seed gene *Lgr6*.

| Seed gene | STRING edges in Metagene | Null expected edges in gene set | Network enrichment | p |
|---|---|---|---|---|
| *Lgr5* | 385 | 94 | 4.096 | 0 |
| *Bmi1* | 204 | 119 | 1.714 | 8.10E-13 |
| *Sox4* | 183 | 112 | 1.634 | 6.60E-10 |
| *Lrig1* | 181 | 86 | 2.105 | 0 |
| *Lgr6* | 218 | 113 | 1.929 | 0 |
| *Sox9* | 271 | 146 | 1.856 | 0 |
| *Sox2* | 106 | 82 | 1.293 | 0.00676 |
| *Prom1* | 303 | 61 | 4.967 | 0 |
| *Foxa1* | 309 | 55 | 5.618 | 0 |
| *Krt15* | 279 | 62 | 4.500 | 0 |
| *Krt19* | 427 | 64 | 6.672 | 0 |
| *Pitx1* | 188 | 63 | 2.984 | 0 |
| *Psca* | 364 | 54 | 6.741 | 0 |
| *Prom2* | 352 | 60 | 5.867 | 0 |
| *Klf5* | 632 | 89 | 7.101 | 0 |
| *Cd44* | 566 | 75 | 7.547 | 0 |

**Table SM1 |** Tumor metagene gene-gene associations indexed by STRING, testing for significant enrichment in network connectivity of carcinoma network metagenes seeded by genes shown in the "Seed gene" column.

**Single-cell network metagene expression stability.** We explored the stability of metagene expression patterns across metagene sizes. Of greatest interest for this study is how an metagene expression pattern changed across single cells as a function of metagene size. We did not attempt a comprehensive quantitative metagene optimization routine in this study since the optimization space is nebulous and very large at this stage of gene network deconvolution. For example, metagenes could be optimized for 1) size within network node number constraints, and the nature of those constraints are specific to particular questions 2) recapitulation of empiric networks, and what empiric networks represent the most functionally relevant eg experimental protein-protein interactions vs. motif binding interactions 3) specificity for certain cell populations, and which populations and how much expression differentiation would be required among them. With this in mind, we examined how broad expression patterns in the single-cell manifold changed as we included an increasingly large number of correlated genes in the network metagene. We were concerned with two qualitative properties of these patterns: 1) sensitivity: sufficiently high metagene expression in a large enough number of cells to detect patterns 2) specificity: expression patterns differentiating among groups of cells. In general, metagene size N=100 balanced these. For example, the *Foxa1* carcinoma metagene was a "lower spike" metagene (it's expression was higher in the lower spike than other papilloma cell populations). Expression patterns began to emerge at relatively large sizes of N=50 and were very specific for the lower spike up to sizes of N=500 (Fig.SM3). On the other hand, for the *Sox2* metagene, expression patterns were immediately apparent at N=10 but specificity rapidly diminished (Fig.SM4). Taken together we considered that metagenes of size N=100 struck the most informative balance between sensitivity and specificity; they encompassed more real network interactions than expected by chance; and occurred near the inflection point of increasing connectivity of protein networks as a function of N.

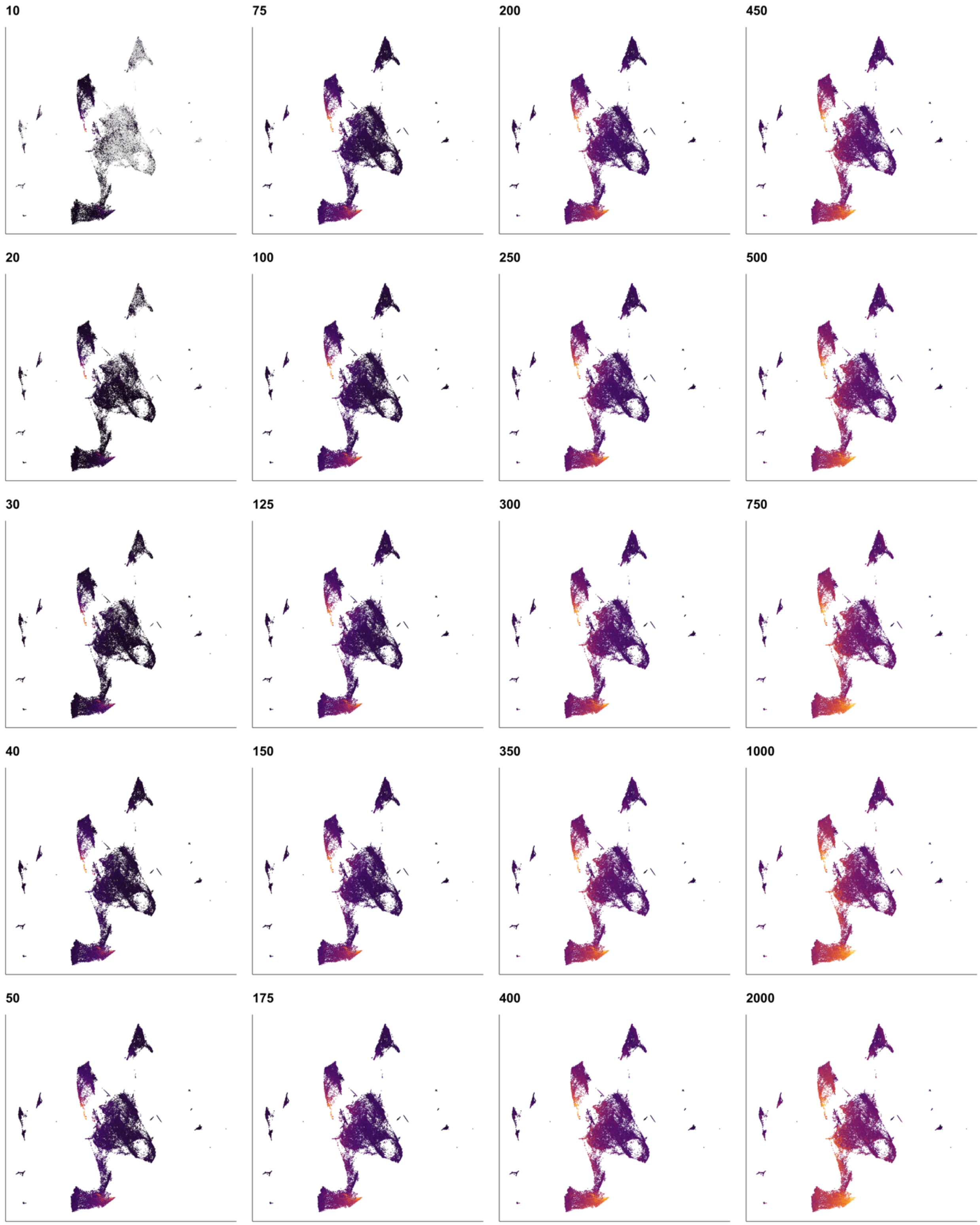


**Fig. SM3 |** *Foxa1* carcinoma metagene expression for a range of metagene sizes from N=10 to N=2000, where N is the number of constituent genes in the metagenes that are most correlated to the seed gene *Foxa1*. Purple is low expression and yellow is high expression. Panel numbers show the number of genes in the network metagene.

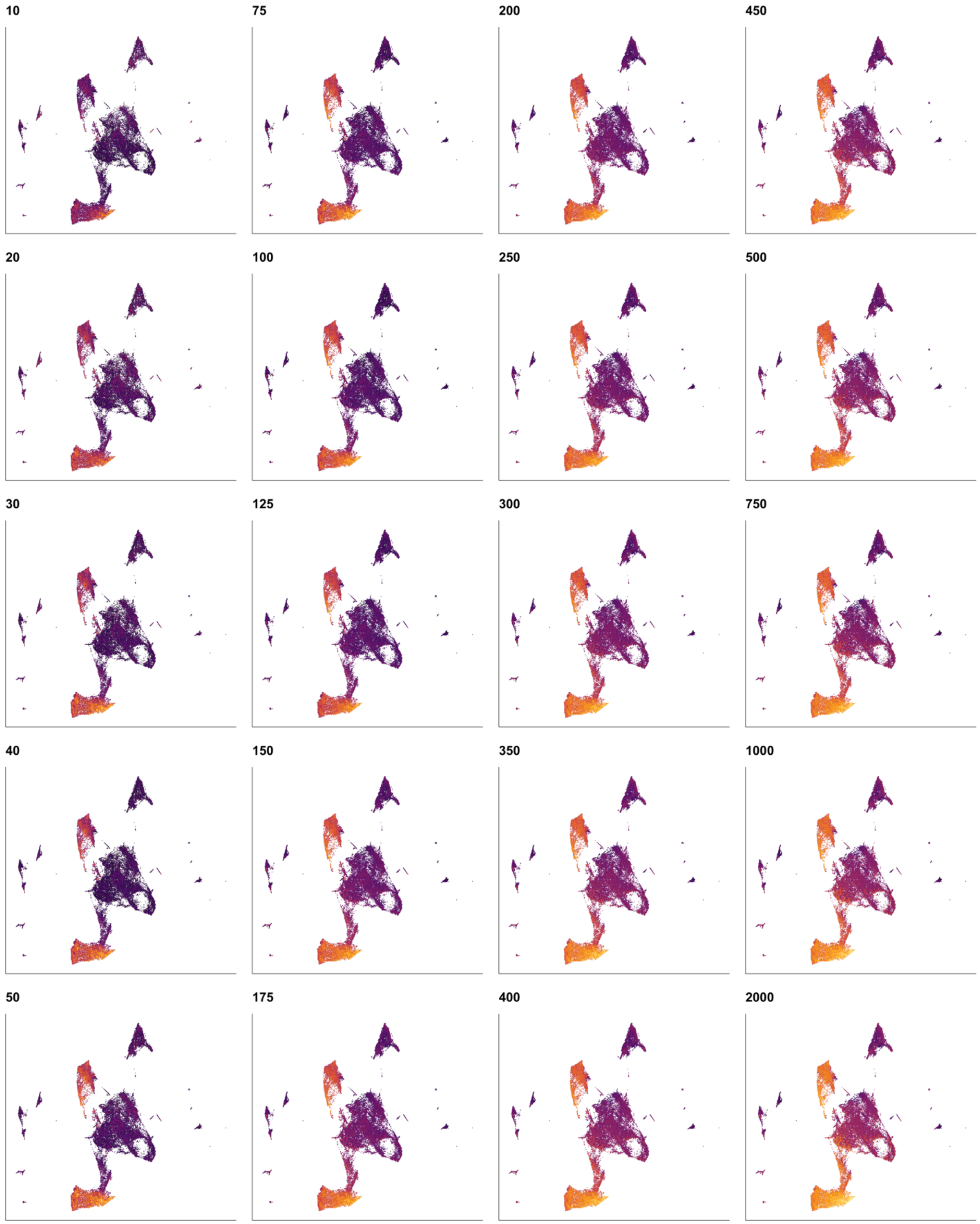


**Fig.SM4 |** *Sox2* tumor metagene expression for a range of metagene sizes from N=10 to N=2000, where N is the number of constituent genes in the metagenes that are most correlated to the seed gene *Sox2*. Purple is low expression and yellow is high expression. Panel numbers show the number of genes in the network metagene.

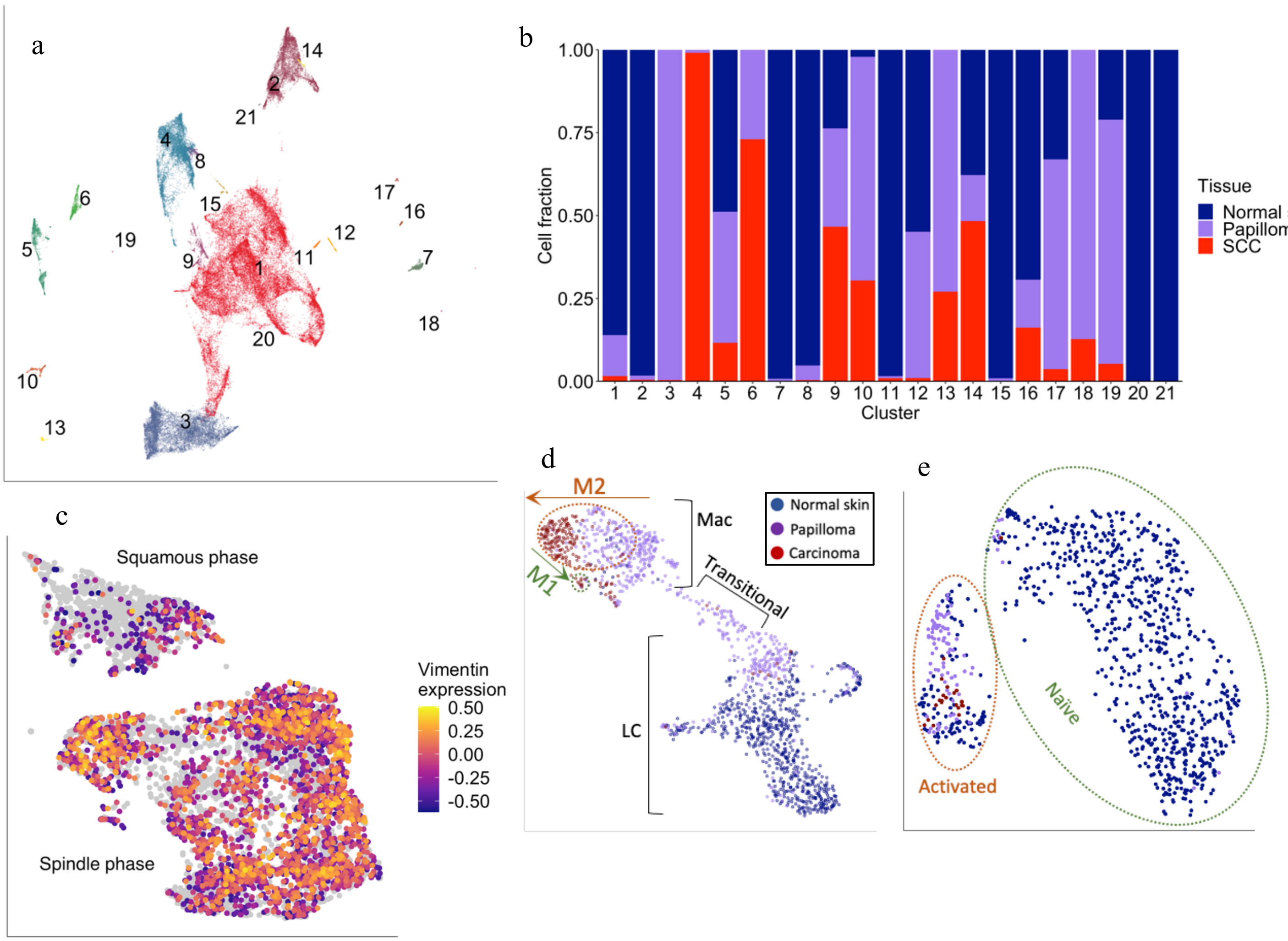


**Fig. S1 a**, Unsupervised Leiden-based cell clusters at default 'partition'-level resolution (k=10, partition q-value=0.05). **b**, Fraction of cells derived from each carcinogenic stage populating unsupervised cell clusters shown in a. **c**, Carcinoma parenchyma cells projected within its own manifold overlain by vimentin expression, showing distinct unsupervised disjunction between the squamous and spindle phases. **d**, Langerhans (LC) and macrophage (Mac) cells projected within their own manifold, and analyzed by cell type, inflammatory markers, and M1/M2 polarization. We consider resident LCs to represent an unactivated cell population in the context of carcinogenesis, marked *by H2-Eb1, H2-Ab1, Cd207, Epcam, Cd24a*, and *Csf1r*. Consistent with this notion, in our carcinogenic series LCs derive entirely from normal skin samples. Macrophages, on the other hand, were entirely drawn from papilloma and carcinoma samples, and were called with markers *Ptprc, Fcgr1, Adgre1, Mertk, Ms4a7, C1qc, Itgam*, and *Cd68*. We tested for the presence of inflammatory macrophages with *Ccrl2, Il1r2, Il1b, Tnf, Dusp2*, and *Kfkbi1*, but no clear axis in the manifold fell out and fewer than 15 cells were defined as inflammatory according to this marker set (see Methods for how cell types were consensually defined). However, we did see strong evidence for an M1/M2 polarization axis. This axis has been criticized as an artifact of *in vitro* challenge by pathogen-associated molecular patterns, classically by the bacterial cell wall molecule lipopolysaccharide (LPS), as well as high influence by tissue microenvironment so that any particular gene set has poor replicability in a different tissue context[1]. Indeed, testing for M1 and M2 macrophages with markers defined by *in vitro* LPS challenge alone[2] did not reveal a clear axis, although it did identify a small subpopulation of

M1 cells. However, testing for the M1/M2 axis using a mouse *in vivo*-derived marker set with both LPS and non-LPS inflammatory challenges[2], revealed a clear gradient across axis 1 of the manifold, and also identified an M2 population that was not apparent using the *in vitro* M2 marker set. Together, this supports the utility of *in vivo* models in understanding monocytic responses to carcinogenesis which are not apparent through classic *in vitro* assays. Finally, transitional cells did not express high levels of either LC or Mac marker sets relative to the LC and Mac populations. Interestingly, these cells were entirely drawn from the papilloma sample, indicating that this premalignant population induces monocytic differentiation and activation along axes ending in macrophage activation. **e**, T-cells projected within their own manifolds. Two groups were clearly present in the data, and we tested for stage-specific differentially expressed genes (DEGs) between these groups. The top DEG with high expression in neoplastic cells was *AW211201*, which has been shown to be a readout of T-cell activation and to promote differentiation of inflammatory T-cells[3]. The second was *Slpi*, which has been shown to be secreted by activated leukocytes in order to protect epithelial tissue from immune protease activity such as leukocyte elastase[4]. Other top DEGs such as *Ctla4, Tnfrsf4*, and *Il74* were consistent with an activated T-cells state and thus we term this cell cluster "activated" and the other cluster "naïve." Importantly, all but 1 carcinoma T-cells were activated as well as the majority of papilloma T-cells, whereas the majority normal skin T-cells were naïve.

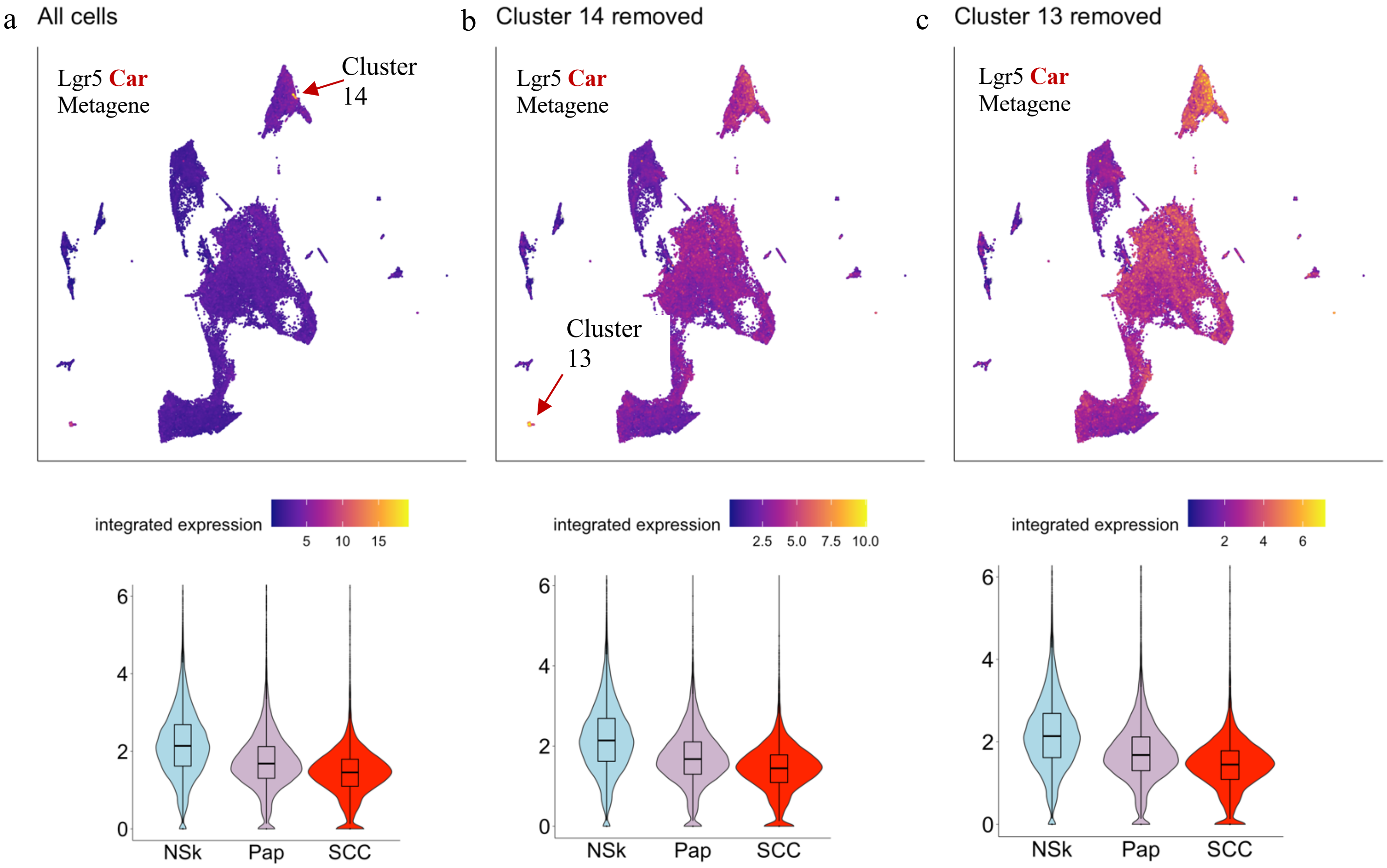


**Fig. S2** Effect of removing rare cell populations (low-resolution cell clusters 13 and 14) with high expression of the *Lgr5* carcinoma metagene on its expression visualization in the UMAP. Cluster 13 consisted of 181 fibroblasts (0.31% of all cells) and cluster 14 consisted of 151 cells (0.26%). If these rare cell populations with high *Lgr5* carcinoma metagene were driving the apparent expression pattern in the UMAP, then we would expect to see a radical transformation in it when these populations were removed. This is not the case. Although it appears that the metagene value is changing in the manifold, it is simply a matter of color rescaling, as shown by the consistent negative trend in the corresponding violin plots. This reveals that the *Lgr5* carcinoma metagene provides a stable negative benchmark for carcinogenesis in our analytical pipeline. Furthermore, the metagene increasingly highlights normal skin follicular cells instead of neoplastic cells, reflecting its known homeostatic function in follicular regeneration. We interpret this to mean that *Lgr5* is not co-opted into pathways important for neoplastic growth relative to *Lgr6*, which is strongly clonogenic in this system and which shows increasing expression across neoplastic progression (Fig. 2c).

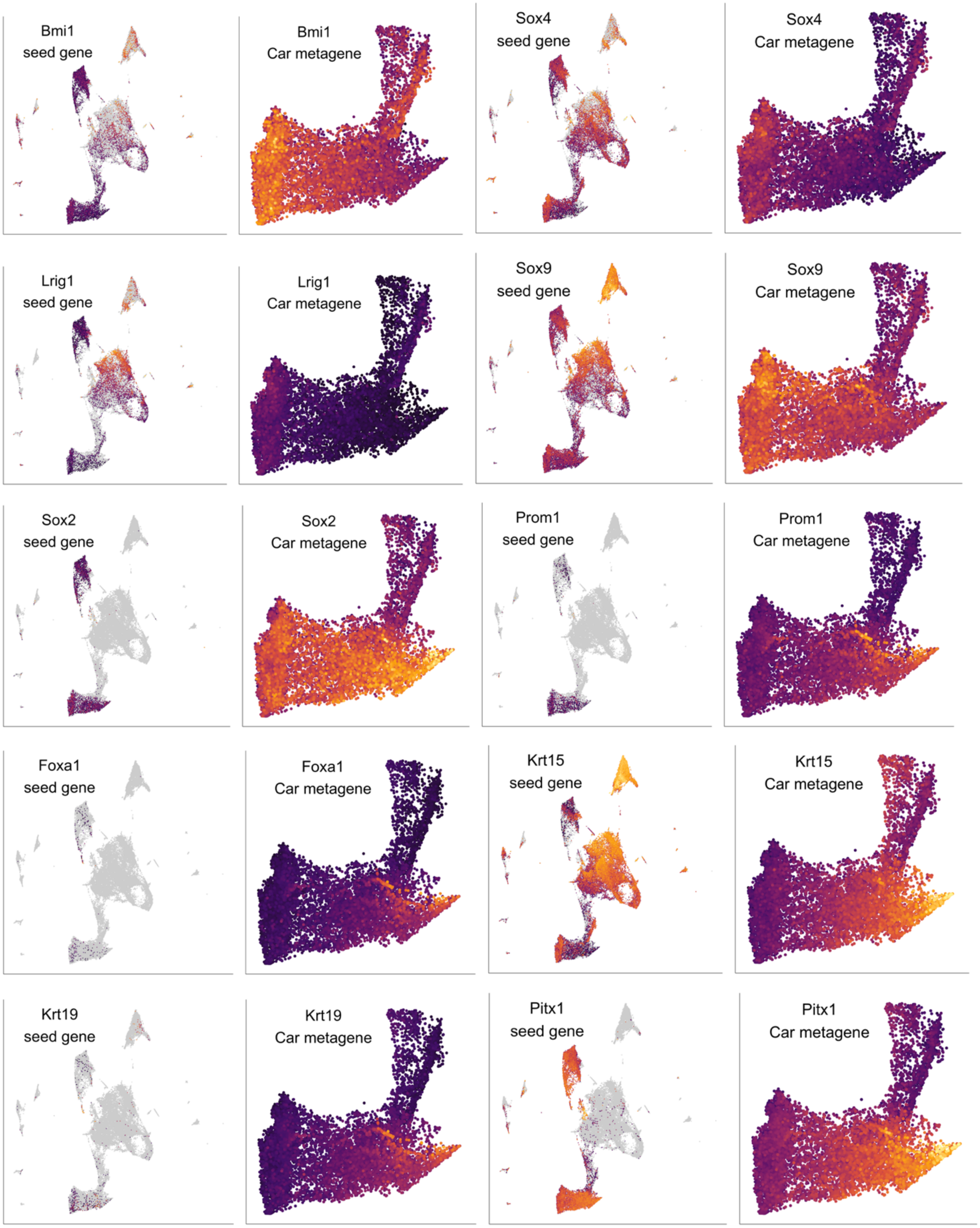
Bmi1
seed gene
Bmi1
Car metagene
Sox4
seed gene
Sox4
Car metagene
Lrig1
seed gene
Lrig1
Car metagene
Sox9
seed gene
Sox9
Car metagene
Sox2
seed gene
Sox2
Car metagene
Prom1
seed gene
Prom1
Car metagene
Foxa1
seed gene
Foxa1
Car metagene
Krt15
seed gene
Krt15
Car metagene
Krt19
seed gene
Krt19
Car metagene
Pitx1
seed gene
Pitx1
Car metagene

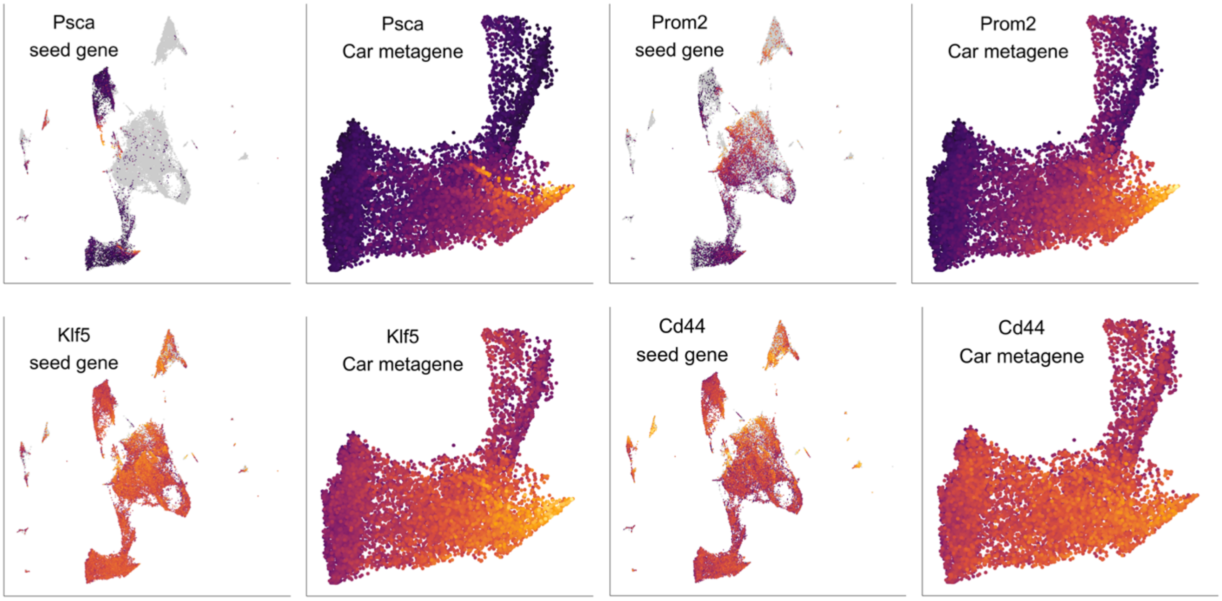


**Fig S3.** Seed gene and carcinoma metagene expression UMAPs for 14 stem-associated genes not shown in the main text. Purple shows low expression and yellow shows high expression. Color gradients are rescaled in each panel.

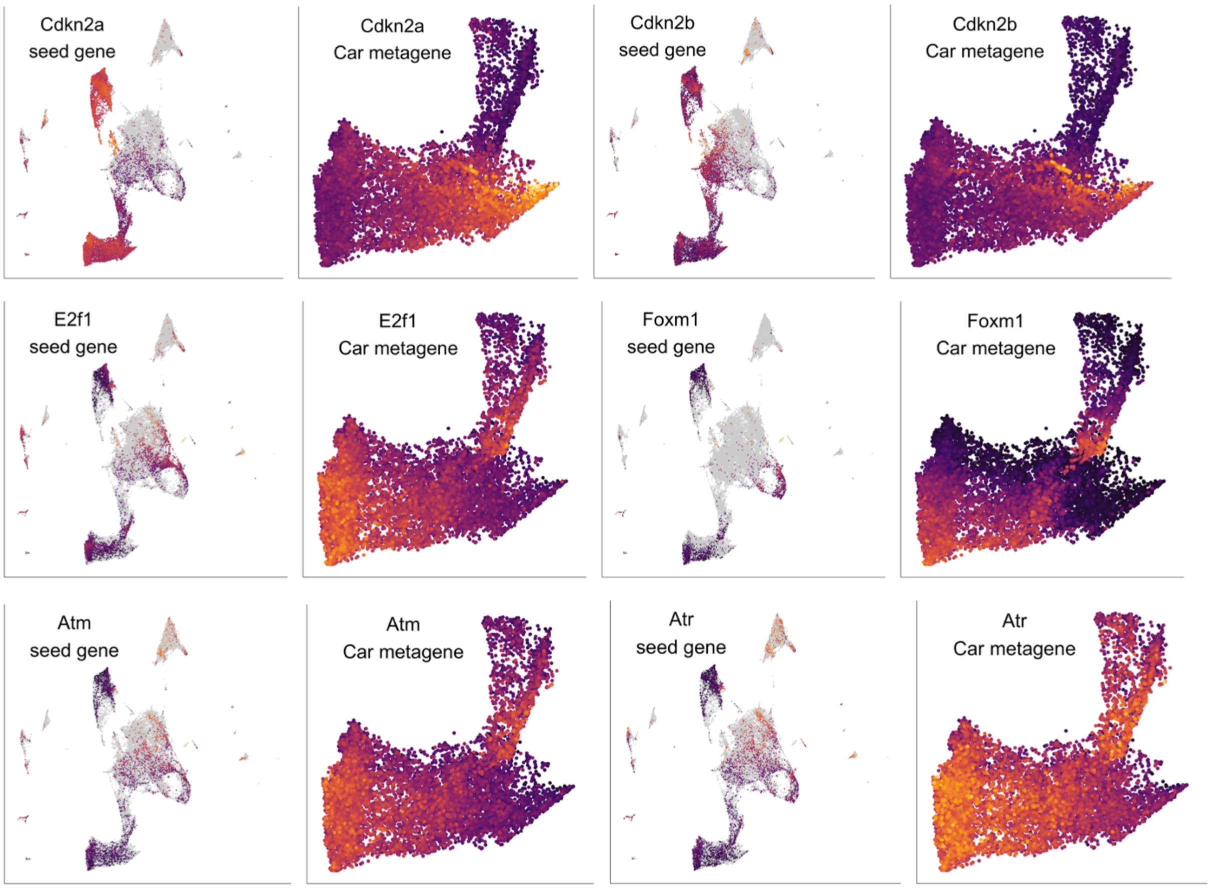


**Fig. S4.** Seed gene and carcinoma metagene expression UMAPs for cell cycle checkpoint genes (*Cdkn2a/b*), cell cycle progression genes (*E2f1* and *Foxm1*), and DNA damage/genomic instability genes (*Atm* and *Atr*). Purple shows low expression and yellow shows high expression. Color gradients are rescaled in each panel.

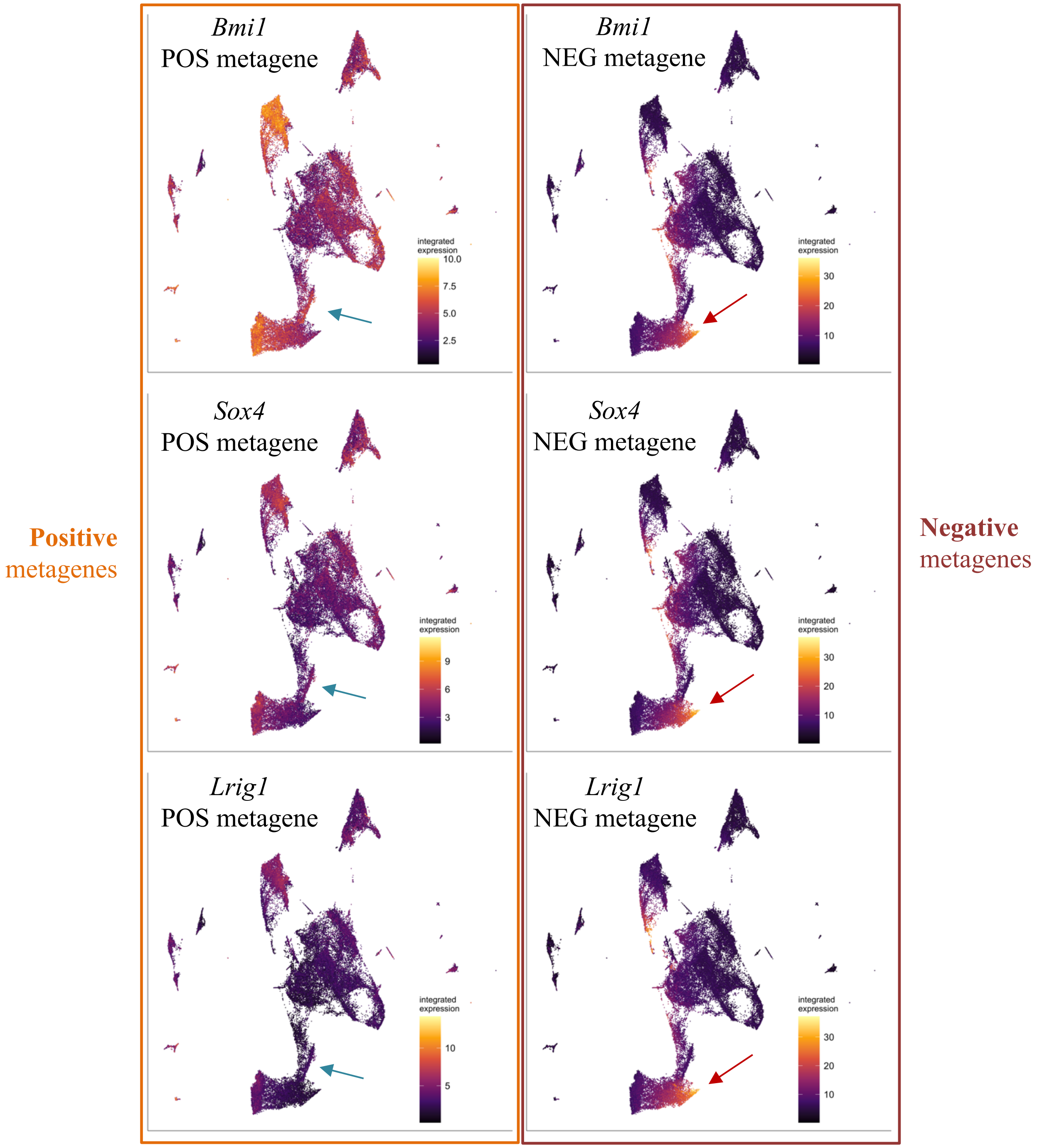


**Fig. S5.** Exemplar positive (POS) and negative (NEG) carcinoma metagene patterns demonstrating that for a given seed gene, its positive metagene is more expressed in one spike and its negative metagene is expressed in the opposite spike of the papilloma parenchyma. A positive carcinoma metagene is defined as the 100 genes most correlated to a seed gene in the bulk-tissue carcinoma samples, and a negative carcinoma metagene is defined as the 100 genes most anticorrelated to a seed gene. For example, for *Bmi1*, its positive metagene is more highly expressed in the US than the LS; however, its negative eigengene is more highly expressed in the LS than the US. This is consistent across spike-specific metagenes, and we take this to reflect the mutual exclusivity of the LS and US network programs that emerge in these alternate cell fates during pre-malignant carcinogenesis.

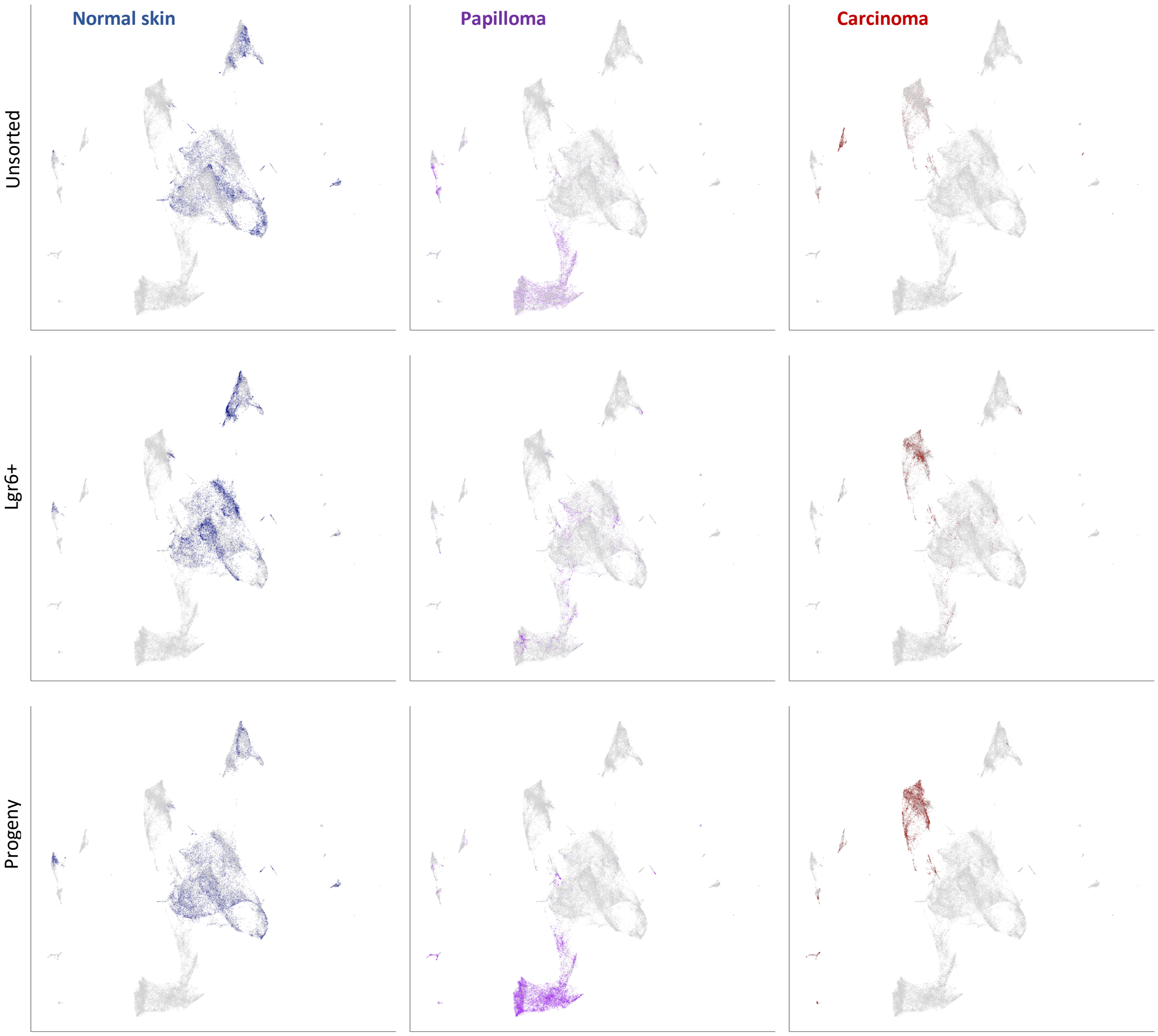


**Fig. S6.** UMAPs showing stage-specific FACS-sorted cells. Columns are malignant stage and rows are FACS sorting status; grey dots are cells that do not fall into a stage×FACS class, whereas colored cells do. This shows that both main neoplastic populations (papilloma and carcinoma parenchyma) consist of a large number of progeny cells, consistent with their being clones initiated by *Lgr6*+ cells of origin.

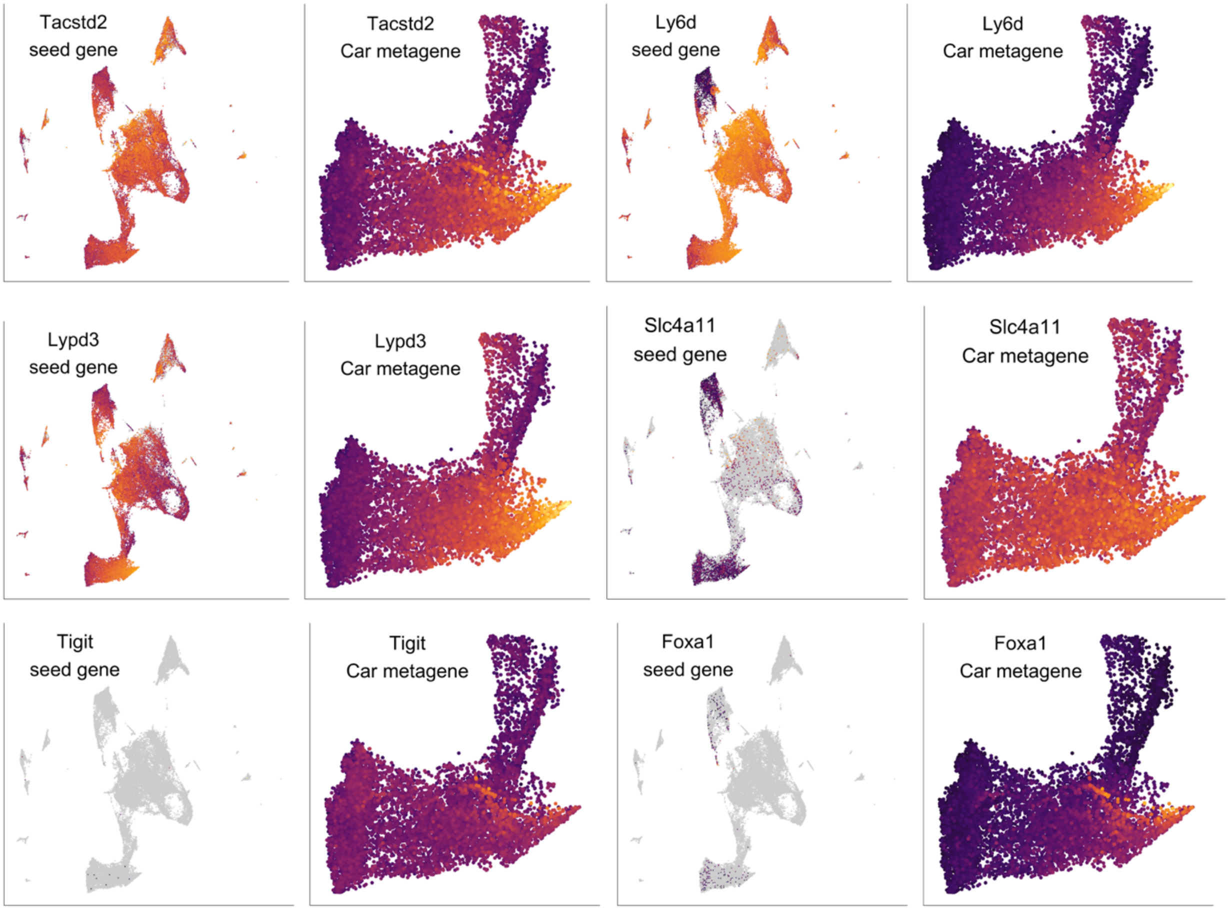


**Fig. S7**. Seed gene and carcinoma metagene expression UMAPs for drug-resistance associated genes from basal cell carcinomas (*Tacstd2, Ly6d,* and *Lypd3*), lung adenocarcinoma (*Slc4a11* and *Tigit*), and prostate adenocarcinoma (*Foxa1*). Purple shows low expression and yellow shows high expression. Color gradients are rescaled in each panel.

| Normal skin *Lgr6* network | | | | Carcinoma *Lgr6* network | | | |
|---|---|---|---|---|---|---|---|
| gene | ρ | gene | ρ | gene | ρ | gene | ρ |
| ***Lgr6*** | 1 | *Cachd1* | 0.767 | ***Lgr6*** | 1 | *Aplp2* | 0.516 |
| *Osbpl5* | 0.868 | *Mtmr11* | 0.766 | *Tnnt2* | 0.722 | *Cd9* | 0.514 |
| *Atp2b4* | 0.852 | *Slc6a2* | 0.763 | *Cnnm4* | 0.711 | *Rai14* | 0.507 |
| *Rapgefl1* | 0.843 | *Cobl* | 0.763 | *Prss12* | 0.672 | *Smurf1* | 0.506 |
| *Phyhip* | 0.843 | *Serpinb7* | 0.762 | *Sox9* | 0.667 | *Nedd9* | 0.506 |
| *Fndc9* | 0.826 | *Gjb5* | 0.761 | *Itga2* | 0.652 | *C1galt1* | 0.505 |
| *Tanc1* | 0.822 | *Col17a1* | 0.761 | *Shf* | 0.644 | *Pmepa1* | 0.502 |
| *Slc24a3* | 0.821 | *Peli3* | 0.76 | *Igsf8* | 0.644 | *Padi3* | 0.501 |
| *Susd2* | 0.818 | *Srcin1* | 0.758 | *Itpr3* | 0.637 | *Gpr25* | 0.501 |
| *Sphk2* | 0.813 | *Eps8l1* | 0.757 | *Nt5e* | 0.613 | *Fermt1* | 0.497 |
| *Rgmb* | 0.813 | *Fam20b* | 0.754 | *Bhlhe41* | 0.612 | *Nav2* | 0.493 |
| *Pou2f3* | 0.808 | *Mark2* | 0.753 | *Fzd7* | 0.6 | *Arhgef3* | 0.49 |
| *Tmem184a* | 0.807 | *Kdm1b* | 0.753 | *Krt18* | 0.599 | *Capns1* | 0.488 |
| *S1pr5* | 0.806 | *Nhsl1* | 0.751 | *Arid5a* | 0.598 | *Foxp4* | 0.488 |
| *Arhgap44* | 0.804 | *Fam129b* | 0.75 | *Tgfb1* | 0.597 | *Ubald2* | 0.485 |
| *Vipr1* | 0.804 | *Il20rb* | 0.748 | *Tes* | 0.592 | *Gm7367* | 0.484 |
| *Gdpd2* | 0.803 | *Elovl1* | 0.748 | *Rap1gap* | 0.59 | *Gm7367* | 0.484 |
| *Axin2* | 0.803 | *Sbsn* | 0.748 | *Add2* | 0.589 | *Ctnna1* | 0.483 |
| *Apcdd1* | 0.8 | *Ano8* | 0.746 | *Sdc1* | 0.587 | *Cd80* | 0.483 |
| *Mrgprb3* | 0.797 | *Sptbn2* | 0.746 | *Jam2* | 0.586 | *Hivep2* | 0.482 |
| *Zbtb7b* | 0.793 | *Ptpn21* | 0.745 | *Gpr39* | 0.58 | *Dyrk3* | 0.481 |
| *Rab3d* | 0.791 | *Efs* | 0.744 | *B4galnt1* | 0.577 | *Rab6a* | 0.48 |
| *4930412O13Rik* | 0.791 | *Gas2l1* | 0.744 | *D8Ertd82e* | 0.571 | *Runx2* | 0.48 |
| *Cldn23* | 0.791 | *Cyp4f39* | 0.743 | *Plekhg3* | 0.571 | *Iqgap1* | 0.479 |
| *Lrp4* | 0.788 | *Efnb2* | 0.743 | *Lamc2* | 0.571 | *Ggct* | 0.479 |
| *Jag2* | 0.787 | *Klhl17* | 0.742 | *Socs2* | 0.559 | *Lrch4* | 0.478 |
| *C2cd2* | 0.787 | *Mesp2* | 0.742 | *Ramp1* | 0.557 | *Skil* | 0.477 |
| *Pdlim2* | 0.786 | *Cdc42bpg* | 0.741 | *Etv6* | 0.556 | *Frrs1* | 0.476 |
| *Wnt7b* | 0.785 | *Scel* | 0.741 | *Lamb3* | 0.546 | *Slc20a2* | 0.473 |
| *Sh3d19* | 0.784 | *Lamb3* | 0.74 | *Egr3* | 0.539 | *Ncmap* | 0.471 |
| *Ascl2* | 0.783 | *Ikzf2* | 0.74 | *Dusp6* | 0.539 | *Reep3* | 0.47 |
| *Krt15* | 0.782 | *Tacstd2* | 0.74 | *Foxp1* | 0.536 | *Nadsyn1* | 0.469 |
| *Ahnak2* | 0.781 | *Il22ra1* | 0.739 | *Dnaja4* | 0.535 | *Bhlhe40* | 0.468 |
| *Efna3* | 0.781 | *Kdm2b* | 0.739 | *Tnfrsf12a* | 0.535 | *Gprc5a* | 0.467 |
| *Mink1* | 0.781 | *Bag4* | 0.738 | *Pip4k2c* | 0.534 | *Slc39a4* | 0.467 |
| *Il17rc* | 0.78 | *Pdzd2* | 0.738 | *Rnf181* | 0.532 | *Hes6* | 0.466 |
| *Camsap3* | 0.779 | *Clstn1* | 0.738 | *Ano1* | 0.532 | *Acsl3* | 0.465 |
| *Krt77* | 0.777 | *Adcy1* | 0.736 | *A130010J15Rik* | 0.531 | *Erc1* | 0.465 |
| *Hdac5* | 0.777 | *Mboat2* | 0.735 | *Il18r1* | 0.531 | *Lama5* | 0.464 |
| *Lce1m* | 0.777 | *Sh3bp1* | 0.734 | *Chst2* | 0.529 | *Ephb4* | 0.463 |
| *Il20rb* | 0.774 | *Fut2* | 0.734 | *Jag1* | 0.528 | *Rmnd5b* | 0.462 |
| *Pik3c2g* | 0.774 | *Eppk1* | 0.733 | *1600029D21Rik* | 0.525 | *Plekha1* | 0.462 |
| *Net1* | 0.773 | *Ly6g6e* | 0.733 | *Tnfsf9* | 0.524 | *Bcr* | 0.462 |
| *Gata3* | 0.772 | *Ptgs1* | 0.733 | *Src* | 0.524 | *Zbtb18* | 0.462 |
| *Il34* | 0.772 | *Abcc1* | 0.731 | *Runx1* | 0.524 | *Shq1* | 0.46 |
| *Tjp3* | 0.771 | *Rxra* | 0.731 | *Rhbdd1* | 0.521 | *Scyl2* | 0.46 |
| *Ppara* | 0.77 | *Slc6a9* | 0.731 | *Smad7* | 0.521 | *Padi4* | 0.46 |
| *Itpr3* | 0.77 | *Ddr1* | 0.729 | *Itga6* | 0.518 | *Scin* | 0.46 |
| *Il20rb* | 0.769 | *Dgcr6* | 0.728 | *Socs2* | 0.517 | *Dhcr7* | 0.459 |
| *Arvcf* | 0.767 | *Trim29* | 0.728 | *Slco2a1* | 0.517 | *Atp10d* | 0.459 |

**Table S1.** Seed-specific gene networks and their edge weights (Spearman rank correlation coefficients [ρ]) correlating to the seed gene *Lgr6*, which together comprise the 100-N first-degree *Lgr6* normal skin and carcinoma networks. Highlighted cells are mentioned in the main text.

| GO term | LS | Expected | p | Contributing genes |
|---|---|---|---|---|
| skin development | 22 | 3.33 | 0.01119 | *St14, Foxq1, Ppl, Cldn4, Evpl, Scel, Anxa1, Grhl3, Sprr3, Tgm1, Barx2, Celsr1, Dsp, Prss8, Tmem79, Grhl1, Grhl2, Itgb4, Jup, Slc27a4, Sprr1a, Tmprss13* |
| wound healing | 15 | 3.93 | 0.00014 | *Ppl, Erbb2, Evpl, Anxa1, Grhl3, Wnt4, Ceacam1, Celsr1, Dsp, Duox1, Klf5, Sox15, Anxa8, Cd44, Rab27a* |
| keratinocyte differentiation | 12 | 1.57 | 0.00814 | *St14, Ppl, Evpl, Scel, Anxa1, Sprr3, Tgm1, Dsp, Tmem79, Grhl1, Grhl2, Sprr1a* |
| positive regulation of cell motility | 10 | 6.56 | 0.00123 | *Rab25, S100a14, Grb7, Anxa1, Cldn7, Duox1, Duoxa2, Fam83h, Ptprz1, Sh3rf2* |
| negative regulation of cell-cell adhesion | 8 | 2.23 | 0.01031 | *Erbb2, Spint2, Anxa1, Arg1, Ceacam1, Gm9573, Cd44, Cdh1* |
| epithelium migration | 8 | 3.29 | 0.01187 | *Rab25, Anxa1, Tacstd2, Ceacam1, Marveld3, Fat2, Grhl2, Jup* |
| sensory perception of sound | 6 | 1.54 | 0.00449 | *Slc9a3r1, Slc52a3, Cdh1, Jag2, Marveld2, Pgap1* |
| cell-matrix adhesion | 6 | 2.35 | 0.00829 | *Wnt4, Itgb4, Itgb6, Jup, L1cam, Lypd3* |
| epithelial cell morphogenesis | 6 | 0.52 | 0.01192 | *St14, Rab25, Spint2, Cdh1, Cldn3, Grhl2* |
| negative regulation of JNK cascade | 3 | 0.44 | 0.00929 | *Ceacam1, Marveld3, Sh3rf2* |
| cuticle development | 2 | 0.02 | 0.00015 | *Duox1, Tmem79* |
| positive regulation of hydrogen peroxide biosynthetic process | 2 | 0.04 | 0.00043 | *Duoxa1, Duoxa2* |
| positive regulation of neutrophil apoptotic process | 2 | 0.06 | 0.00142 | *Anxa1, Cd44* |

**Table S2.** Gene ontology (GO) enrichment for the 223 genes shared among the 10 Car eigengenes with high expression in the lower spike (cluster 3 seed genes). GO term refers to the biological annotation attached to the term; annotated refers to the number of genes across the single-cell transcriptome that belong to the GO term; LS refers to the number within 223 genes that belong to the GO term; expected refers to the number expected given the size of the GO term if there were no enrichment; p refers to the significance of the Fisher exact test for enrichment; genes in term show the 223 genes that are annotated by the particular GO term.
